# A naturally occurring urinary collagen type I alpha 1-derived peptide inhibits collagen type I-induced endothelial cell migration at physiological concentrations

**DOI:** 10.1101/2025.07.08.663687

**Authors:** Hanne Devos, Ioanna K. Mina, Foteini Paradeisi, Manousos Makridakis, Aggeliki Tserga, Marika Mokou, Jerome Zoidakis, Harald Mischak, Antonia Vlahou, Agnieszka Latosinska, Maria Roubelakis

**Author notes:** Present address of Hanne Devos: Department of Cardiovascular Disease, Nordic Bioscience A/S, DK-2730 Herlev, Denmark. Correspondence: Maria Roubelakis or, Agnieszka Latosinska.

## Abstract

Collagen type I (COL(I)) is a key component of the extracellular matrix (ECM) and is involved in cell signalling and migration through cell receptors. Collagen degradation produces bioactive peptides (matrikines), which influence cellular processes. In this study, we investigated the biological effects of nine most abundant, naturally occurring urinary COL(I)-derived peptides on human endothelial cells at physiological concentrations, using cell migration assays, mass spectrometry-based proteomics, flow cytometry, and AlphaFold3. While none of the peptides significantly altered endothelial migration by themselves at physiological concentrations, full-length COL(I) increased cell migration, which was inhibited by Peptide 1 (^229^NGDDGEAGKPGRPGERGPpGp^249^). This peptide uniquely contains the DGEA and GRPGER motifs, interacting with integrin α2β1. Flow cytometry confirmed the presence of integrin α2β1 on human endothelial cells, and AlphaFold3 modelling predicted an interaction between Peptide 1 and integrin α2. Mass spectrometry-based proteomics investigating signalling pathways revealed that COL(I) triggered phosphorylation events linked to integrin α2β1 activation and cell migration, which were absent in COL(I) plus peptide 1-treated cells. These findings identify Peptide 1 as a biologically active COL(I)-derived peptide at a physiological concentration capable of modulating collagen-induced cell migration, and provide a foundation for further investigation into its mechanisms of action and role in urine excretion.

## 1. Introduction

The extracellular matrix (ECM) is a specialized three-dimensional network, that plays a critical role in cell proliferation, survival, migration and differentiation, through interactions with various cell receptors [1]. Among the ECM’s components, collagen type I (COL(I)) is one of the most abundant proteins in both the ECM and the human body [2]. Dysregulation of COL(I) has been implicated in numerous diseases [3], most notably fibrotic conditions characterized by an imbalance between collagen synthesis and degradation [2]. COL(I) is composed of two alpha 1 chains (COLα1 (I)) and one alpha 2 chain (COLα2 (I)). Upon maturation, COL(I) molecules are exported to the ECM, where they form cross-links – primarily between hydroxylated proline and lysine residues – to create robust collagen fibrils. These fibrils interact with cell surface receptors, such as integrin α1β1 and integrin α2β1 [2,4]. Activation of these receptors by collagen initiates intracellular signalling cascades, including mobilization of Src kinase and activation of Focal Adhesion Kinase (FAK) [5,6]. Downstream pathways involving key signalling proteins like Extracellular-signal Regulated Kinase (ERK) and Akt are subsequently activated [6–13], promoting processes such as cell migration, proliferation and angiogenesis [5,7,8,10,14,15].

Matrix-metalloproteinases (MMPs) [16] and cathepsins [17] are prominently involved in the degradation of ECM proteins. This degradation generates peptides, some of which exhibit biological activity and are termed “matrikines”. These matrikines can circulate in blood and are frequently detectable in urine [18,19]. Their identification is facilitated by technologies such as capillary electrophoresis coupled with mass spectrometry (CE-MS), a robust and reproducible high-throughput method [20]. Biological activities of collagen-derived matrikines were reported in the context of ECM deposition, collagen synthesis, chemo-attraction, proliferation, and inflammation [18]. Specific motifs on these peptides, such as the DGEA motif, have been extensively studied. Synthetic peptides containing the DGEA motif interact with integrin α2β1 [21], activating pathways involving Signal Transducer and Activator of Transcription 6 (STAT6) [22], c Jun N-terminal kinase (Jnk) [23], Akt [24], and calcium mobilization through inositol 1,4,5-triphosphate (IP3) and phospholipase C (PLC) [25–27]. Functionally, DGEA has been shown to inhibit endothelial cell migration [28], suppress angiogenesis [29], influence macrophage polarization toward an M2 phenotype [22,30], and induce osteogenic differentiation [23,31,32] and cell adhesion [21,26,31]. Other motifs, such as GUXGEZ (where U is usually hydrophobic, X usually hydroxyproline, and Z is often arginine), also interact with integrin receptors [33]. Prominent examples include GFOGER, GROGER and GLOGER motifs (with ‘O’ denoting hydroxyproline), which bind to integrin α1β1 (GFOGER, GROGER) and α2β1 (GFOGER, GLOGER, GROGER) [33,34]. While the signalling pathways for GROGER– and GLOGER-motifs are not well described, GFOGER-containing peptides are known to activate FAK phosphorylation [35–37] and Akt signalling via PI3K [38]. The minimally functional GER motif has been linked to inhibition of angiogenesis [39,40]. Moreover, the PGP motif has been reported to interact with C-X-C chemokine receptor 1/2 (CXCR1/2) [41–43], activating pathways such as GRP-Rac1, p21-Activated Kinase (PAK) and ERK [44], while reducing Akt phosphorylation [45]. PGP also induces inflammatory responses, including neutrophil chemotaxis [41–43] and cytokine secretion [46], and promotes endothelial progenitor cell migration, proliferation and neovascularization [47]. Additionally, the RGD motif, which interacts with integrin α5β1 and integrin αvβ3 after collagen denaturation, has similar biological effects [48–51]. Comparable activities have been reported for matrikines derived from other types of collagen [52]. However, most studies on matrikines and collagen activity rely on short synthetic peptides at high concentrations, rather than on the naturally occurring collagen-derived peptides at physiological levels. Naturally occurring collagen-derived peptides have been associated with conditions such as kidney, cardiovascular and liver diseases, and have been proposed as biomarkers for these disorders [53–56]. These diseases often involve compromised endothelial function [57,58]. Understanding how abundant collagen peptides influence endothelial cells is therefore biologically relevant and may reveal mechanisms through which these peptides impact vascular health and disease progression.

Previous research using CE-MS/MS to analyse peptides in plasma and urine from 22 subjects highlighted a significantly higher abundance of COLα1 (I)-derived peptides in urine compared to plasma, with strong correlations observed among their relative abundances in both samples [59]. In contrast, non-collagen-derived peptides showed poor correlations between their urine and plasma abundances [59]. This discrepancy may be due to active reabsorption mechanisms in kidney tubules, which likely prevents most plasma peptides from being detectable in urine [60]. However, collagen-derived peptides may be exempt from this reabsorption process, potentially due to their potential biological activity.

Based on the above reports, we investigated whether abundant, COLα1 (I)-derived peptides [59] influence endothelial cell migration at physiological concentrations, as reported for full length COL(I) [7,15,61–63]. In a second step, potential molecular mechanisms for an observed impact on endothelial cells are investigated. The methodology and an overview of results are summarized in **Figure 1**.

**Figure 1:**
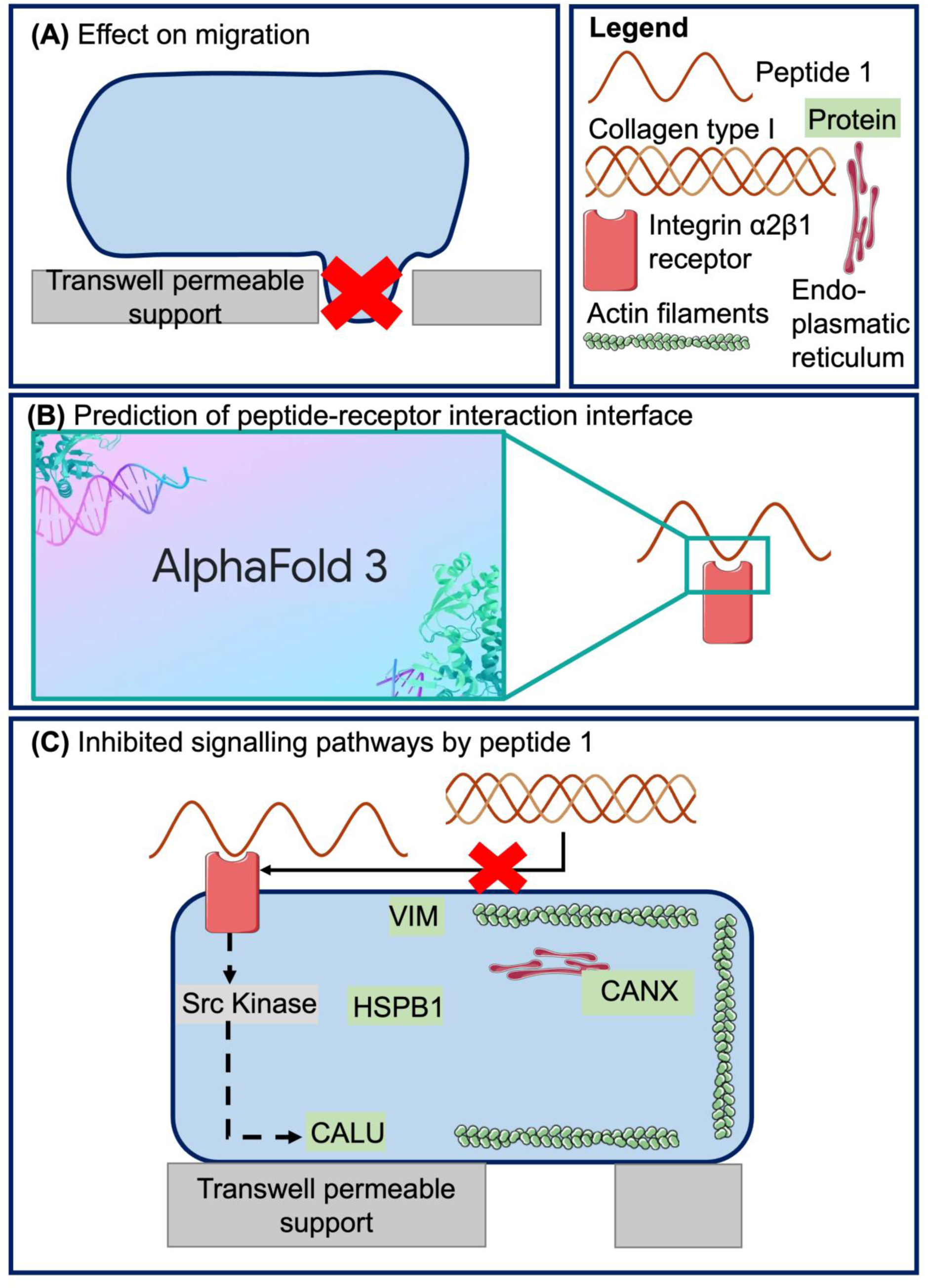
Overview of the methodology and results. First, we investigated the effect of the peptides on cell migration (A). Next, we confirmed the presence of the suggested receptor (integrin α2β1) based on the literature and modelled the interaction of the biologically active Peptide 1 with the receptor using AlphaFold 3 (B). Lastly, we investigated the signalling pathways inhibited by the biologically active peptide (C).

## 2. Results

### 2.1. Shortlisting of peptides likely to have biological activity based on abundance

We hypothesized that collagen-derived peptides are excluded from tubular reabsorption due to potential biological effects on endothelial cells. Accordingly, we assumed that the more abundant a collagen-derived peptide is in urine, the more likely it is to have biological activity. Based on the findings of Magalhães et al. [59], we shortlisted nine highly abundant, non-overlapping COLα1 (I)-derived peptides identified in urine.

A closer examination of these peptide sequences (**Table 1**) revealed the prominent presence of the PGP motif previously reported to interact with cell-surface receptors [41–47,64], in seven out of the nine peptides. One of these peptides, in addition to the PGP motif [41–47,64], also contained the DGEA motif [21–32] and the GRPGER motif [33,39,40,65]). No known motifs were identified in the remaining two peptides. In line with previous analyses of the urinary peptidome of COLα1 (I) [66], the PGP motif, which appears 51 times in the mature COLα1 (I) sequence, was frequently detected, while peptides containing the DGEA or GRPGER motifs-appearing twice and once respectively in the mature COLα1 (I) [67] were much less frequent [66].

**Table 1:**
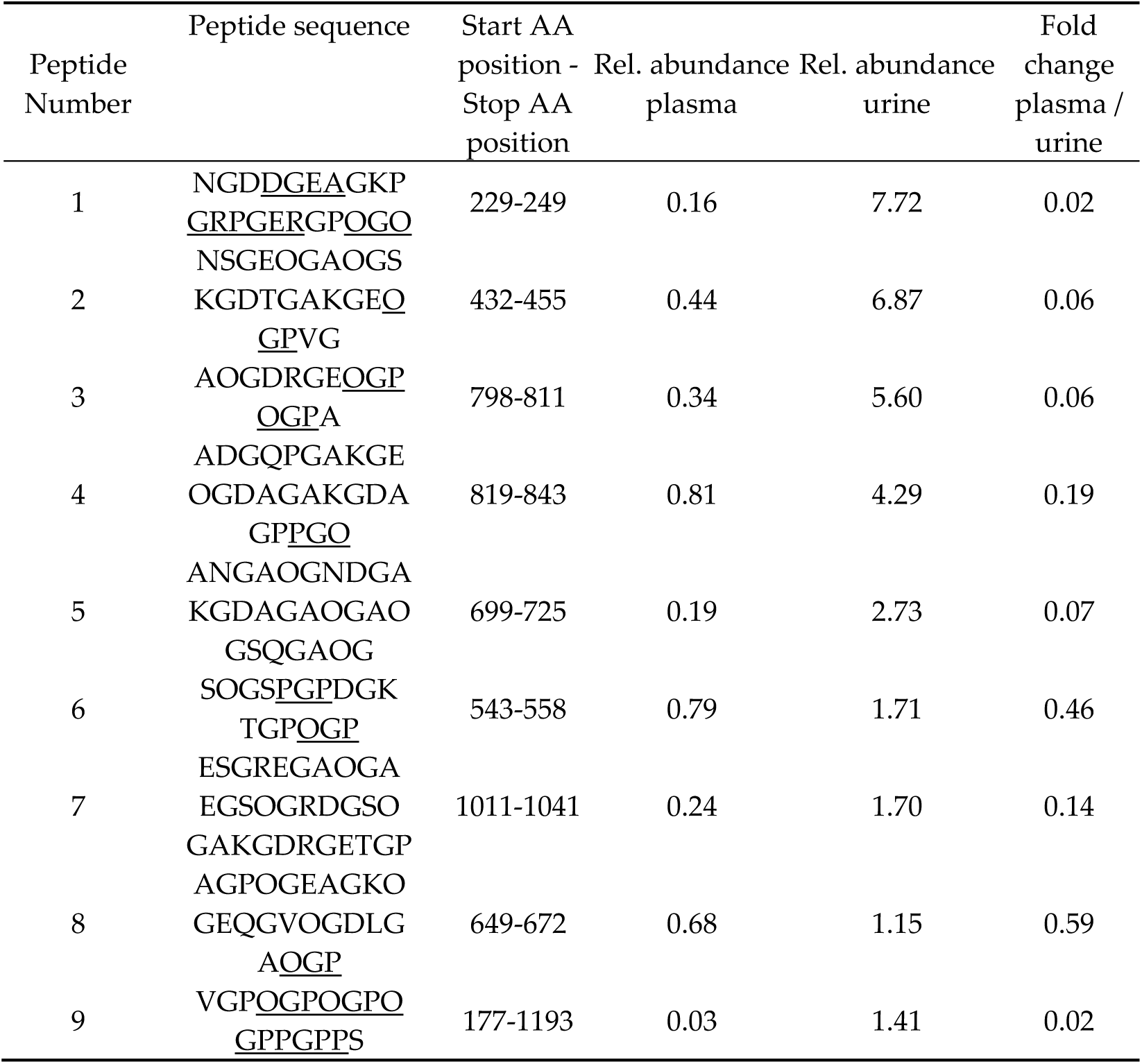
Overview of the shortlisted peptides investigated in this study. The table presents their names, sequences (with bold indicating motifs known to interact with cell receptors and ‘O’ denoting hydroxyproline), as well as the start and stop positions on the collagen type I alpha 1 chain. Additionally, the relative abundance in plasma and urine, along with the fold change in plasma over urine, as calculated in Magalhães et al. [59] are shown, rounded to two decimals.

### 2.2. Inhibition of collagen-induced migration by a naturally occuring urinary peptide

The selected naturally occurring COL(I)-derived peptides were tested at physiological concentrations (100 nM, as further explained in section 4.2) in a migration assay using Human Umbilical Vein Endothelial Cells (HUVEC) cells. Full-length COL(I) was applied as a control. COL(I) treatment doubled endothelial cell migration compared to the negative control (p-value < 0.001, mean number of migrated cells 30.58 for COL(I)-treated vs. 17.02 for negative control). Surprisingly, none of the nine peptides significantly affected endothelial migration (**Supplementary** Figure 1). Mean migrated cell counts and standard deviations for all treatments are reported in **Appendix B, Table B1**, a comparison of the number of migrated cells for peptide-treated cells is shown in **Supplementary Figure S1**, with representative microscope images in **Supplementary Figure S2**. However, one peptide (Peptide 1 (^229^NGD**DGEA**GKP**GRPGER**GP**pGp**^249^, containing the DGEA, PGP, and GRPGER motifs), significantly inhibited collagen-induced migration at physiological concentrations, reducing cell migration by 50% (p-value < 0.001; mean number of migrated cells 30.58 for COL(I)-treated vs. 15.92 for COL(I) + peptide 1-treated cells) (**Figure 2**). No effect on COL(I)-induced migration was observed for the other peptides at physiological concentrations.

**Figure 2:**
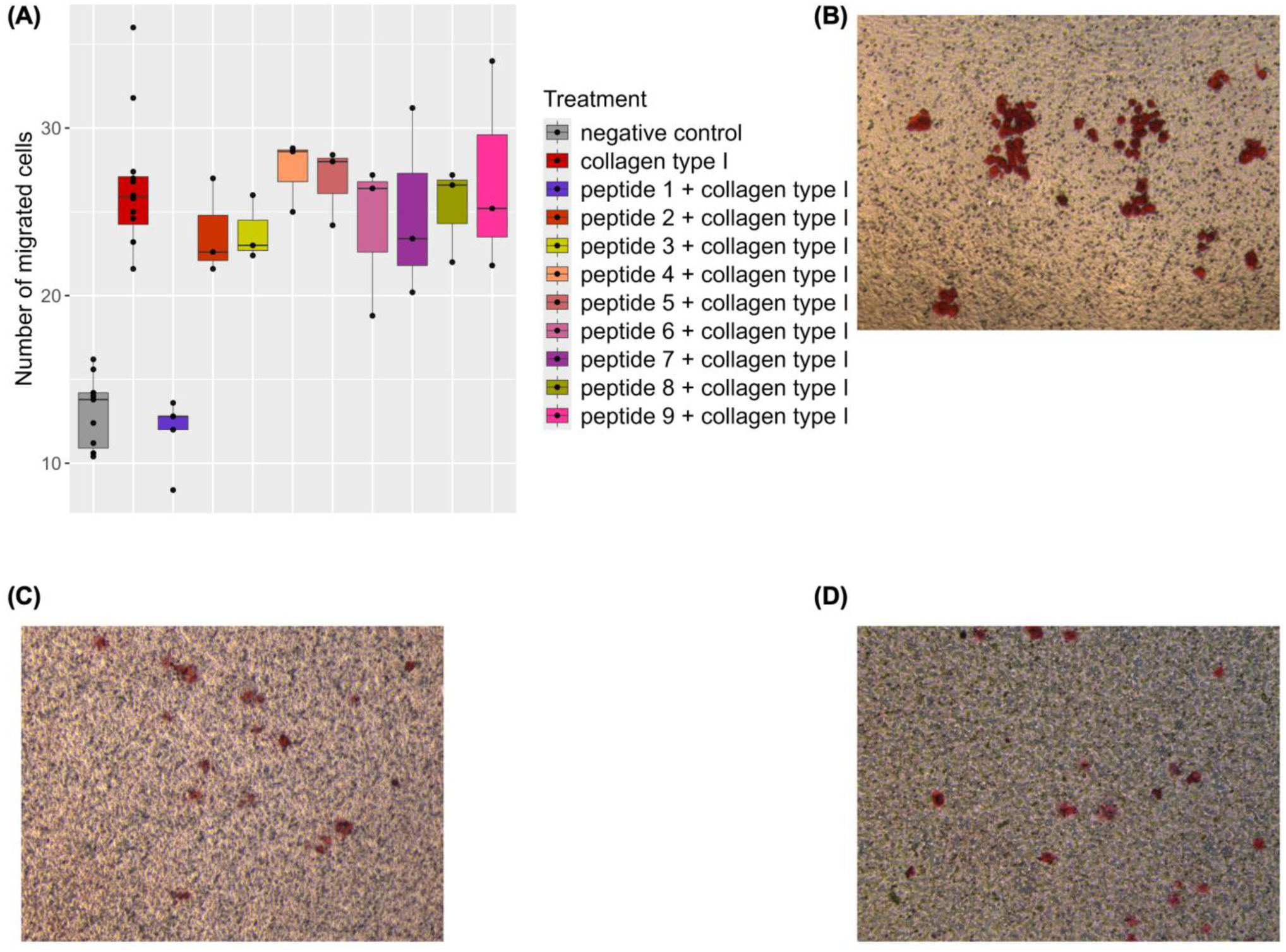
(A) boxplots of the average number of migrated cells per biological replicate, per treatment condition. Results from 3 replicates are shown, except for the negative control (n=11), COL(I)-treated cells (n=12), and peptide 1 + COL(I) (n=5). Data analysis utilized a negative binomial generalized linear model, followed by post-hoc Welch two-sample t-tests (p < 0.05). B-D) Representative microscopic images of the migration experiment for COL(I) (B), negative control (C), and Peptide 1 + COL(I) (D). Images taken at 10X magnification.

### 2.3. Modelling peptide 1 and integrin α2β1 receptor interaction with AlphaFold3

Flow cytometry analysis confirmed the expression of integrin α2β1 receptors on HU-VEC cells (**Figure 3A**), confirming the proteomic identification of integrin α2 and β1 in all samples (**Supplementary File**, tab integrin receptors). To explore peptide-receptor interactions, AlphaFold 3 [68] was employed to model the interaction of Peptide 1 with integrin α2β1. The model consistently predicted proximity of Peptide 1 residues ^242^GER^244^ (GRPGER motif) with integrin α2 residues ^183^NS^155^, and Peptide 1 residue ^245^G with residue ^219^D of integrin α2 (**Figure 3B**). Integrin receptor positions were based on UniProtKB annotations after pro-peptide elimination [69,70].

**Figure 3:**
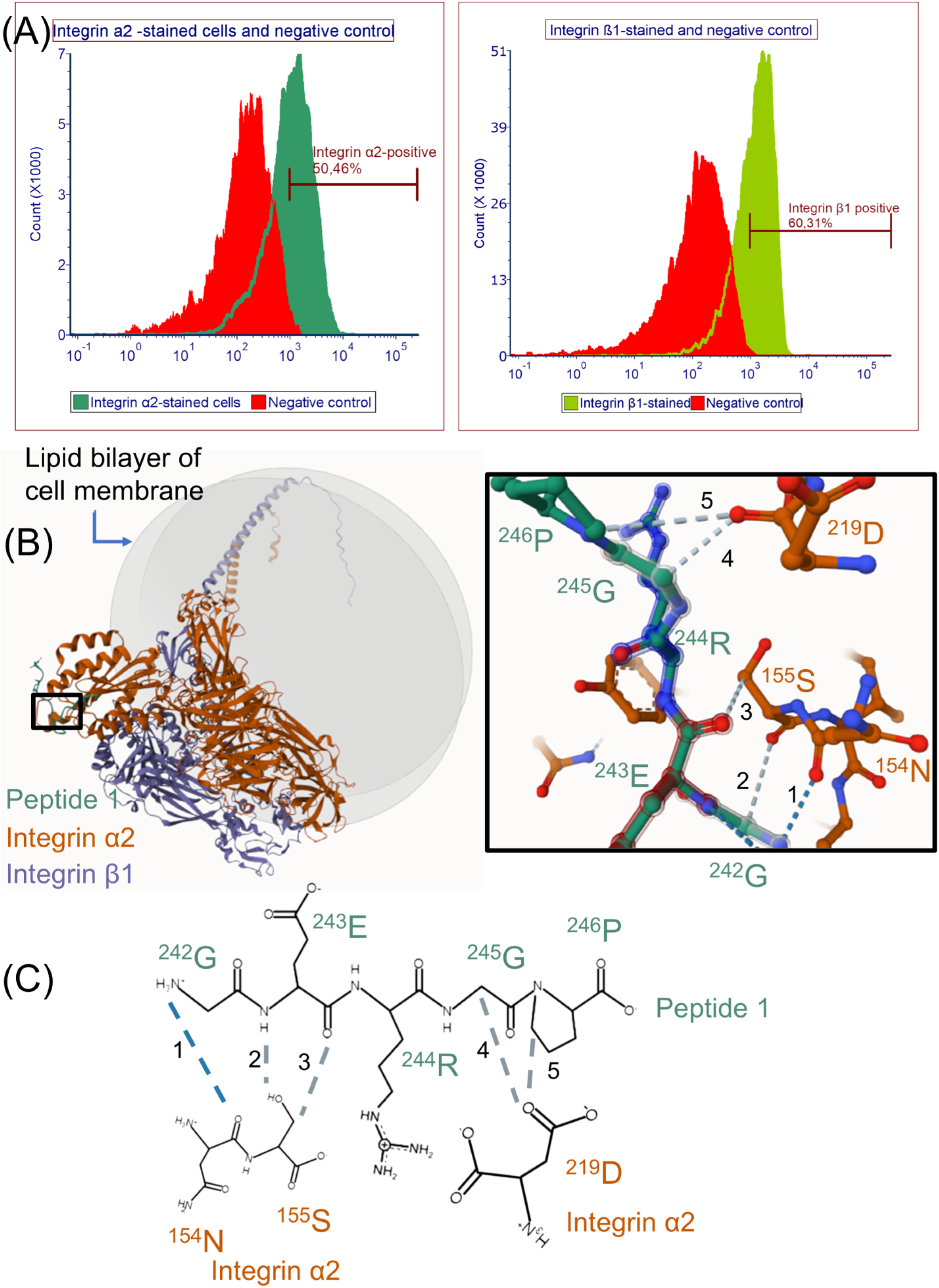
(A) Flow cytometry analysis of the integrin α2β1 receptor. The plot on the left shows that the integrin α2 receptor is present on more than half of the cells, while the plot on the right shows a similar distribution for integrin β1. (B) The left panel shows the AlphaFold 3 predictions of the integrin receptor–Peptide 1 complex. The integrin α2 subunit is represented in orange, integrin β1 in purple, Peptide 1 in green, and the position of the cell membrane relative to the predicted peptide-receptor complex is indicated by the grey double disk (representing the lipid bilayer). Residues predicted to be within 8Å of each other are considered interacting, as defined by AlphaFold3 and widely accepted in the field (see for example [68,71–73]), and are visualised in more detail with Mol* software. The predicted integrin α2 receptor-Peptide 1 interaction is enlarged on the right (other residues were excluded from the right-handed image to increase clarity). Dashed lines indicate predicted interactions, with grey lines indicating weak and blue lines indicating strong interactions. 1: Peptide 1 residue ^242^G and ^154^N on integrin α2, 2: Peptide 1 residue ^242^G with ^155^S on integrin α2, 3: Peptide 1 residue ^243^E with integrin α2 residue ^155^S, 4: Peptide 1 residue ^245^G with integrin α2 residue ^219^D and 5: Peptide 1 residue ^246^P with integrin α2 residue ^219^D. Remaining bonds are between integrin α2 residues internally or peptide 1 internally. (C) Schematic representation of the residues relevant to the predicted peptide 1-integrin α2β1 interaction are shown. Their names are coloured according to the molecule they belong to (orange for integrin α2 and green for peptide 1), dashed lines indicate predicted interactions, with grey lines indicating weak and blue lines indicating strong interactions. Numeration of the bonds is the same as for panel (B). Images in part (C) were drawn using MolView (https://app.molview.com/).

### 2.4. Investigation of potentially affected signaling pathways

We explored signalling pathways reported to be activated by COL(I), and the effects of Peptide 1 at physiological concentrations. Initial Western Blot analyses (**Supplementary Figure S3**) suggested activation of pERK, a known downstream effector of collagen I stimulation [6–13]), however, the results were not consistent, and the observed potential impact was low. Therefore, mass-spectrometry was employed to identify phospho-peptides in HUVEC cells treated with COL(I) alone or in combination with Peptide 1.

Twenty-one phospho-peptides were identified with high-confidence in the COL(I) or COL(I) + Peptide 1-treated samples (**Figure 4A; Supplementary File**, tab differential abundance testing). Six of these phospho-peptides exhibited significant differences in abundance between Peptide 1 + COL(I)-treated samples and COL(I)-treated samples (**Figure 4A**), five downregulated, one upregulated. Among these, four phospho-peptides (originating from CALU, CANX, HSPB1 and VIM) showed significantly decreased abundance in the Peptide 1 + COL(I)-treated samples as compared to the COL(I)-treated samples across at least two of the time points tested (7min, 30min, 60min) (**Figure 4B**). **Table 2** summarizes the phospho-peptides with significant differential abundance between the two tested conditions (Peptide 1 + COL(I) versus COL(I)).

**Figure 4:**
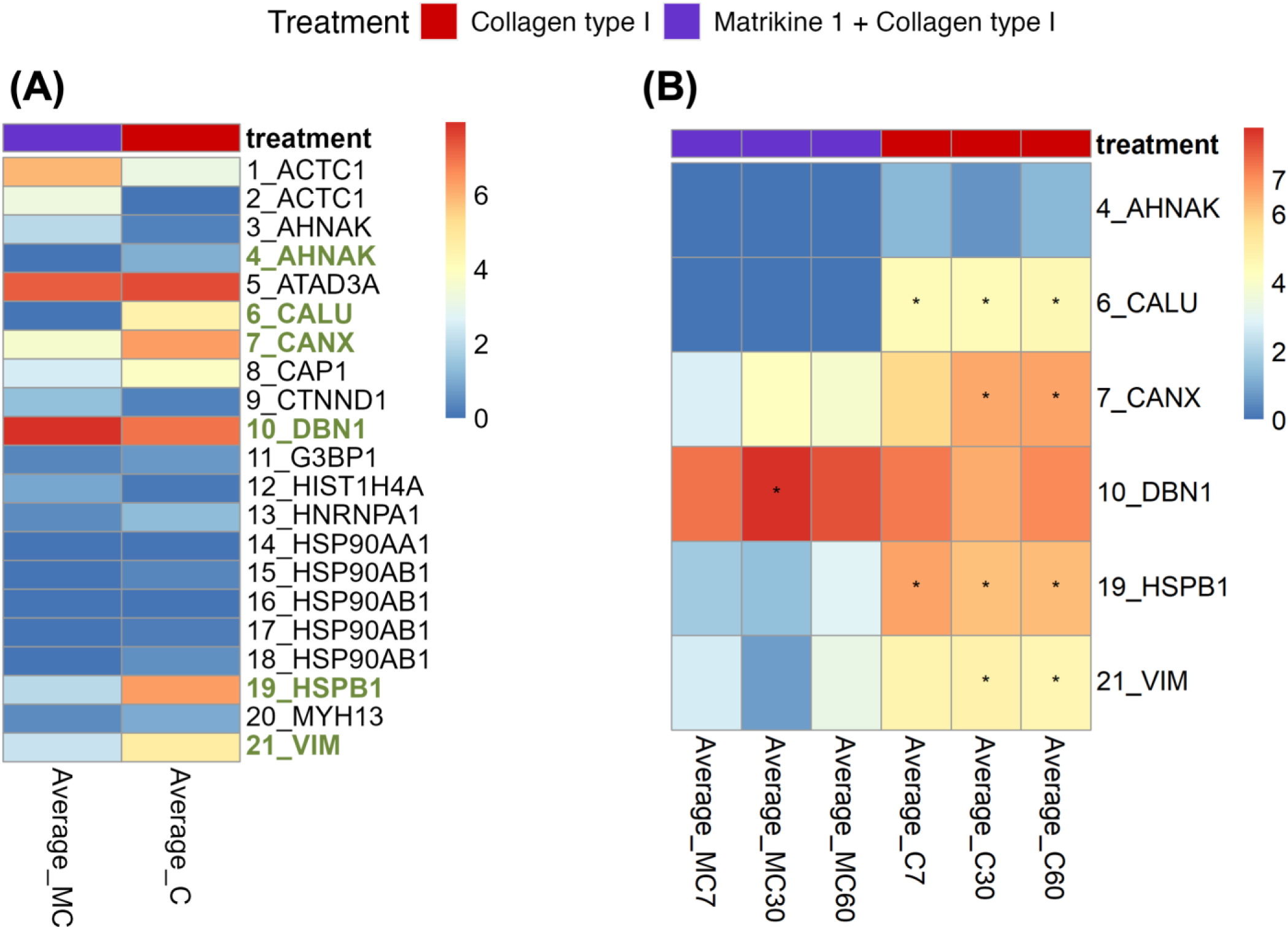
(A) Average abundance per treatment, with the proteins of which peptides display significantly differential abundance (after Benjamini-Hochberg correction) between treatments highlighted in bold and green. Data from twelve replicates are shown. (B) Breakdown of the average abundance per time point for the significant phospho-peptides shown in panel (A). Asterisks denote significant differences in abundance at specific time points and are displayed in the column corresponding to their highest abundance. For each time point and per treatment, four replicates were evaluated.

**Table 2:**
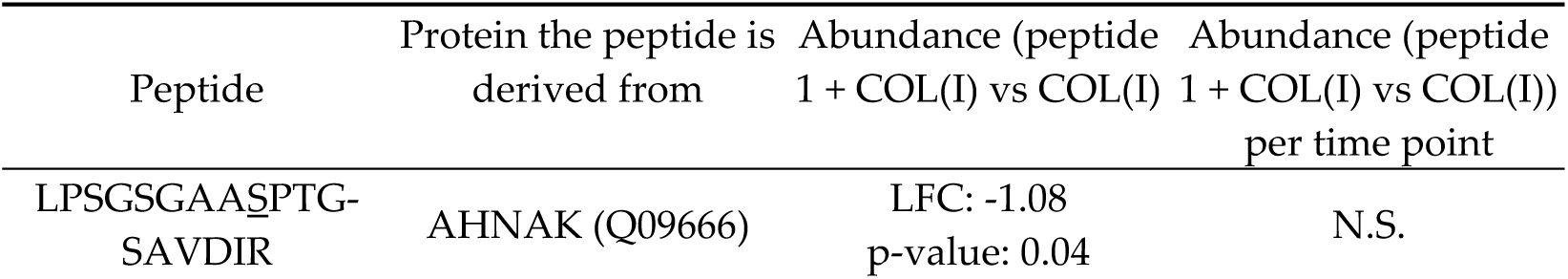
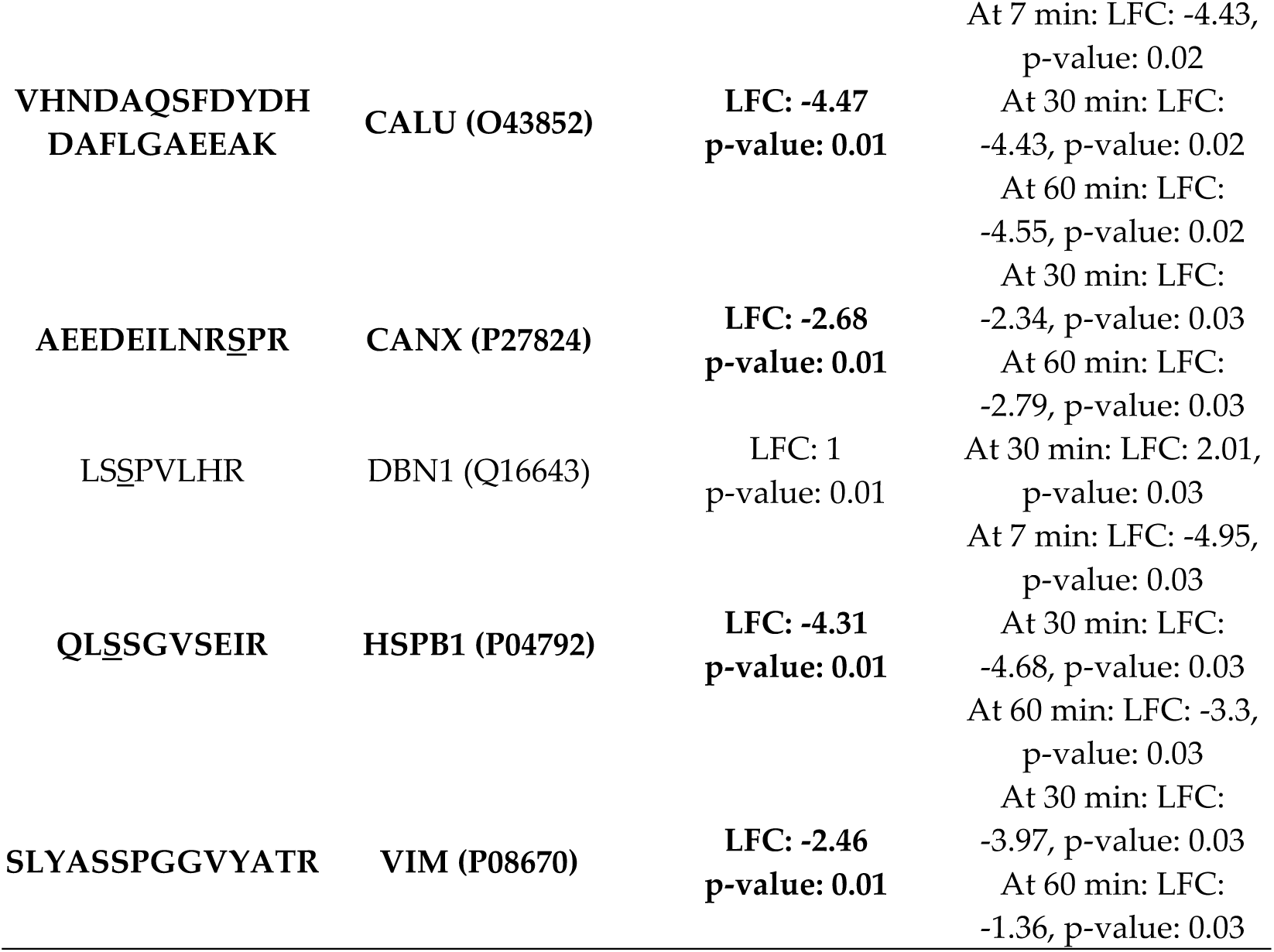
Overview of the detected phospho-peptides with statistically significant differences in their abundance between Peptide 1 + COL(I)-treated samples and COL(I)-treated samples. Phosphorylated residues in the peptide sequences are underlined (for CALU and VIM, the phospho-site could not be determined). Statistically significant differences in abundance were assessed using the Wilcoxon rank test, first for each treatment and then for each treatment and time point. Abbreviations: LFC: log fold change; AHNAK: Neuroblast differentiation-associated protein AHNAK; CALU: Calumenin; CANX: Calnexin; DBN1: drebrin; HSPB1: heat shock protein beta 1; VIM: vimentin. Peptides showing significant changes in at least two out of three time points are shown in bold. Means and standard deviation for each analysis are provided in Supplementary File, tab “differential abundance testing”.

## 3. Discussion

COLα1 (I)-derived peptides have been associated with conditions such as kidney, cardiovascular and liver diseases, and have been proposed as biomarkers for these disorders [53–56]. We investigated the hypothesis that urinary COLα1 (I)-derived peptides may exert biological effects on endothelial cells, based on prior findings [59] suggesting that these peptides, unlike those derived from other proteins, may not undergo active tubular reabsorption, of yet unknown mechanism [59]. In this study, we examined nine abundant, non-overlapping urinary COLα1 (I)-derived peptides identified in Magalhães et al. [59]. Among these, only one peptide, Peptide 1 (^229^NGDDGEAGKPGRPGERGPpGp^249^) demonstrated a significant effect in attenuating collagen-induced endothelial migration at a physiological concentration. Notably, Peptide 1 alone, like all other peptides tested, had no measurable effect at the tested concentration (**Figure 2, Supplementary Figure S1**). This peptide is one of the peptides part of biomarker panels for kidney, cardiovascular and liver diseases [53–56]. Interestingly, these diseases often involve compromised endothelial function [57,58], and more specifically, impaired endothelial cell migration [74,75].

Collagen-induced endothelial migration is well documented [61,63,76], and synthetic peptides containing sequences from COLα1 (I), such as DGEA [28,29], PGP [47], and GER-containing peptides [39,40], have been shown to influence migration. However, these synthetic peptides may not accurately represent the behaviour of the naturally occurring peptides, as they frequently were used in much higher than the physiological concentration (even at micromolar levels), and/or only very short peptides encompassing these 3-6 amino acids were investigated, while in general much larger peptides, containing the respective motifs, are observed in vivo. To our knowledge, this study is the first to report a biological function for a naturally occurring urinary COLα1 (I)-derived peptide at physiological concentrations, representing an advancement beyond existing studies that used high concentration of non-naturally occuring peptides, and thus providing findings likely more reflective of physiological conditions. Other studies have highlighted the biological activity of naturally occurring peptides in plasma, for example a study by Lindsey et al. [77] identified, through mass spectrometry, a naturally occurring COLα1 (I)-derived peptide in the plasma of myocardial infarction patients, which was generated by cleavage of the C-terminal region of COLα1 (I), 37 amino acids upstream of the collagen C-telopeptide. Lower levels of this peptide in the plasma were correlated with reduced cardiac function. The “activity peptide” (p1158/59, sequence AcRTGDSGPAGPPGPPG-NH2), enhanced fibroblast migration and wound closure rates in vitro, while in vivo, in mouse models, it improved cardiac function by attenuating increases in left ventricular end-diastolic and end-systolic diameters post-myocardial infarction. Despite its promising cardioprotective effects in a mouse model, the receptor, functional motif, and signalling pathways involved remain unidentified [77]. These findings, together with our results, highlight the therapeutic potential of naturally occurring peptides.

### 3.1 Biological activity of peptide 1 and suggested interaction with integrin α2β1

Our findings suggest that Peptide 1, which uniquely contains DGEA and GRPGER motifs among the investigated peptides, may interact with the integrin α2β1 receptor. This receptor has been previously implicated in collagen interactions, particularly through motifs like GFOGER [33,34]. Previous research highlights PGP-induced endothelial cell migration [47], nevertheless it should be noted that the peptide used in these earlier experiments was acetylated. Furthermore, PGP acts through interaction with CXCR [41–43]. Considering HUVECs only have low expression of CXCR1 [78], this may explain the discrepancy with our results (i.e. no impact of PGP-containing peptides on migration), and underscores the need for further testing the biological impact of such peptides on multiple additional cell lines at physiological concentrations. We confirmed the presence of the integrin α2β1 receptor on HUVEC cells and used AlphaFold 3 to predict proximity between Peptide 1 and receptor residues. Predicted proximity included residues ^242^GER^244^ (part of the GRPGER motif) proximal to ^154^N [79,80], ^155^S [80], and residue ^245^G proximal to ^219^D [79,80] of integrin α2. These residues are critical for collagen binding to integrin α2β1 [81], as confirmed by site-directed mutagenesis and adhesion assays in human platelets [82]. It is important to note that, while AlphaFold3 predictions support our hypothesis, direct experimental confirmation of Peptide 1 binding to integrin α2β1 is still required.

### 3.2 Investigation of signaling pathways

To understand the mechanisms underlying the effects of Peptide 1, we investigated its impact on signalling pathways activated by COL(I). Initial attempts using Western blot analysis failed to detect significant effects (i.e. significant differential phosphorylation), likely due to insufficient sensitivity for subtle changes. Moreover, while based on previous literature phospho-Akt and phospho-ERK are likely to be involved, based on the signalling pathways activated by the investigated motifs (cf. Introduction) [24,38,44,45], such studies frequently employed higher concentrations of regularly short peptides (shorter than the used naturally-occurring peptides), which may alter the observed effects on cell signalling (eg an effect may be less pronounced or delayed). Therefore, we opted to employ mass spectrometry-based phospho-proteomics which has increased sensitivity and is an untargeted approach. The spectra of the suspected phospho-peptides were manually evaluated for credibility, leveraging respective previously published strict identification criteria [83,84]. Using this stringent analysis, six significantly affected phospho-peptides were successfully identified. Four of these phospho-peptides, previously associated with cell migration, were significantly increased in COL(I)-treated samples. This increase was abrogated in samples treated with both COL(I) and Peptide 1 (**Supplementary File**, tab differential abundance testing; **Table 2**).

The literature provided additional insights into these phospho-peptides and their potential roles in migration. Vimentin is phosphorylated at 56S by PAK1, which is required for integrin β1 activation. Inhibition of PAK1 reduces focal adhesion sites and cell extensions, while constitutively active PAK1 enhances them [85]. Heat Shock Protein Beta 1 (Hsp27), phosphorylated at 82S, prevents filament degeneration and promotes polymerization, facilitating cell migration [86,87]. Knockdown of Hsp27 or its upstream kinases inhibits VEGF-induced migration in HUVEC cells [88]. Calumenin, a target of Src kinase [89], is associated with increased cell migration and may act through ERK signalling pathways [90]. Notably, SRC kinase is activated by collagen binding to the integrin α2β1 [5,6]. Phosphorylated calnexin is involved in the folding of newly synthesized N-linked glycoproteins [91], which are increased in the cell membrane of migrating endothelial cells [92]. While these results were carefully examined (both evaluating the spectra and examining their relevance in the literature), and support the results of the migration assays, further validation and also studies investigating the exact signalling pathways triggered by the receptor and explaining the migration read-out are needed.

### 3.3 Proposed model of peptide 1 action

A model summarizing our findings and the literature is presented in **Figure 5**. Briefly, COL(I) activates integrin β1 receptors, a process facilitated by phosphorylated vimentin [85]. This activation triggers downstream signalling, including Src kinase-mediated phosphorylation of calumenin, which supports migration [90]. Phosphorylated Hsp27 contributes to actin filament bundling and filopodia formation, essential for cell migration [86,88,93]. Phosphorylated calnexin aids in the folding of N-linked glycoproteins, which later move to the cell membrane and affect cell adhesion and migration [91,92].

**Figure 5:**
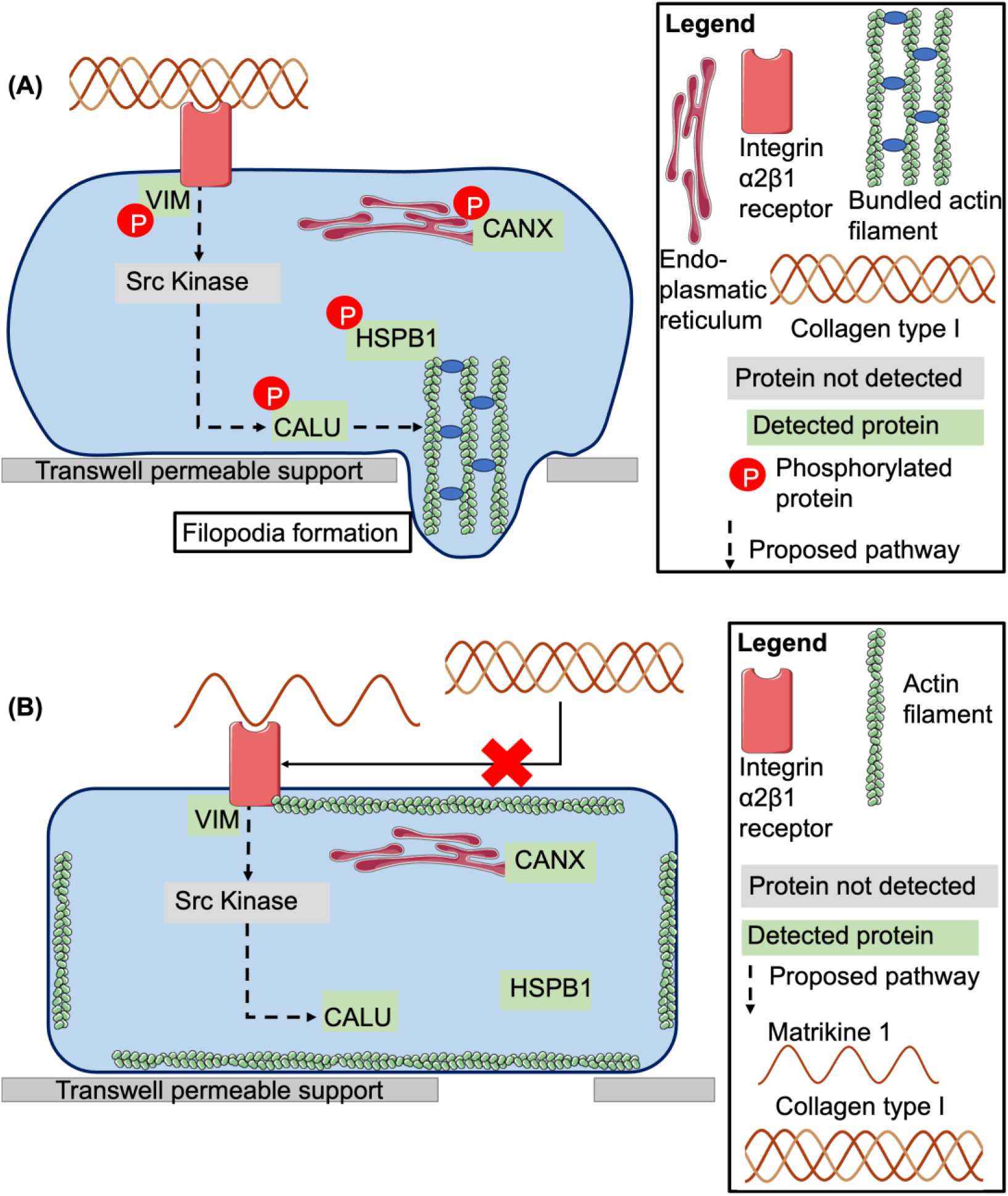
(A) Overview of the proposed signalling pathways activated by COL(I). COL(I) binding to the integrin α2β1 receptor triggers the activation of Src kinase [5,6] (which was not detected in this study), leading to the downstream activation of calumenin (CALU) [90]. Heat shock protein beta 1 (HSPB1) and CALU are involved in the formation of filopodia and subsequent endothelial cell migration [86,88,90,93]. Phosphorylated calnexin aids in the folding of N-linked glycoproteins, which later move to the cell membrane and affect cell adhesion and migration [91,92]. (B) Peptide 1 likely blocks the interaction of COL(I) with integrin α2β1, thereby preventing the activation of the signalling cascade outlined in panel A. Abbreviations: VIM: vimentin, CALU: calumenin, HSPB1: heat shock protein beta 1, DBN1: drebrin.

Peptide 1 appears to reduce the respective phosphorylation and thereby inhibits activation of these pathways by COL(I) at a physiological concentration. A possible mechanistic explanation could be that Peptide 1, by binding to only one integrin α2β1 receptor as a monomer, prevents receptor crosslinking. Mature collagen, as a trimer, would typically crosslink two receptors and so initiate signalling [94]. However, the interaction of peptide 1 with integrin α2β1 and its proposed effect on the receptor crosslinking remains to be experimentally confirmed.

## 4. Materials and Methods

### 4.1 Cell culture

HUVEC were chosen for their fidelity in modelling human endothelial cells and their sensitivity to extracellular matrix (ECM) changes [95]. Pooled HUVECs (Gibco – Thermo Fisher Scientific, C0155C) were cultured at 37°C with 5% CO_2_ under sterile conditions in Endothelial Growth Medium 2 (EGM2; Lonza, CC-3162), with 2% Fetal Bovine Serum (FBS) (Gibco – Thermo Fisher Scientific, 11570506) initially supplemented with 20% FBS until the first passage post-thaw. Thereafter, cells were maintained in EGM2 supplemented with 10% FBS. Experiments were conducted using cells at passage 5 or 6, with the same passage used for biological replicates. Experimental conditions for each assay are described in their respective sections. All cells tested negative for Mycoplasma contamination and were confirmed positive for Cluster of Differentiation 31 (CD31) using immunohistochemistry.

### 4.2 Peptides and collagen type I

Physiological concentration for the peptides was calculated as follows: in a previous study, we assessed the physiological concentrations of the COL1 peptides based on the analysis of a standard urine sample representing normal urine from healthy individuals [96]. This information, together with the average relative abundance of these peptides measured by mass spectrometry in 555 samples from individuals in the general population, was used to estimate the physiological concentrations of the COL1 peptides investigated in this study. For the nine peptides analyzed, the average concentration ranged from 14 nM to 95 nM. Specifically, for matrikine 1, the 95% confidence interval was 15.6 to 127.7 nM. The distribution of matrikine 1 concentrations in the 555 general population samples is presented in **Supplementary** Figure 5. As the concentration ranges varied across peptides, we tested a uniform concentration of 100 nM for all, which falls within the range of physiological concentrations.

Urinary collagen-derived peptides (sequences listed in Table 1, based on [59]), were purchased from Genosphere Biotechnologies (Clamart, France). Peptides were resuspended in cell culture grade water and used at a final concentration of 100 nM in Phosphate Buffered Saline (PBS). Human Collagen Type I (COL(I); Sigma, C7774) was solubilized in 0.5M acetic acid at a concentration of 0.33 mg/ml used at a final concentration of 10 nM (1,4 μg/ml) in PBS.

### 4.3 Migration assay

HUVECs were serum-starved in a 10:1 mix of Endothelial Basal Medium (EBM2; Lonza, CC-3156) and EGM2 (final FBS concentration: 0.2%) for 24 hours before the experiment. Cells were harvested with trypsin-Ethylenediaminetetraacetic acid (EDTA; Gibco, 15400054) and seeded into transwell permeable supports (Costar, 3421) with a 5.0 μm pore polycarbonate membranes at a density at 15,000 cells per insert. Treatments listed in **Appendix B, Table B1** were added to the lower wells (100 nM peptides and 10 nM COL(I), or the same volume of acetic acid dilution in PBS as was added for COL(I)). Migration occurred over 16h [97], after which medium and non-migrated cells were removed. Migrated cells were fixed in 4% paraformaldehyde (PFA) and stained with haematoxylin and eosin. Five representative images per insert were captured at 10X magnification (LEICA DMIRE2 microscope; LAS X software) and analysed using ImageJ’s cell counter plugin (v2.14). Inserts showing damages were excluded from the analysis. Repeat experiments were conducted for treatments with initial promising results (COL(I), Peptide 1 + COL(I) and Peptide 1). Biological replicates are shown in **Appendix B, Table B1**, and each biological replicate has five technical replicates. Data analysis utilized a negative binomial generalized linear model, followed by post-hoc Welch two-sample t-tests (p < 0.05).

### 4.4 Flow cytometry analysis

Medium was removed and cells were washed with PBS and harvested using 0.05% EDTA (Gibco, 15576028). After centrifugation (200g), cells were resuspended in growth medium, washed again, and centrifuged at 360g, followed by washing in PBS + 0.5% Bovine Serum Albumin (BSA; PanReacAppliChem, A1391). For each FACS tube, 5×105 cells were prepared. Cells were incubated with Fc block (Miltenyi Biotec, 130-059-901) for 10 minutes at room temperature to reduce nonspecific binding, then stained with primary antibodies for integrin α2 (CD49b, – FITC–conjugated; BD Biosciences Cat# 555498, RRID:AB_395888) and integrin β1 (CD29; BD Biosciences Cat# 556048, RRID:AB_396319)) in PBS + 0.5% BSA for 1 hour at 37°C in the dark. Cells were washed using PBS + 0.5% BSA, and cells stained with integrin β1 were treated with a secondary antibody (Rabbit Anti-Mouse IgG, RPE-conjugated; Agilent Cat# R043901, RRID:AB_579537), and washed as before. All cells were incubated with 7-Aminoactinomycin D (7-AAD, BioLegend, 640922) for live-dead staining. Calibration and quality control of the BD FACS Celesta instrument were performed before analysis. Data was analysed using FCS Express v7.24.0024 (https://denovosoftware.com/), selecting the cell population, gating out dead cells and comparing double-positive stained cells to isotype controls.

### 4.5 Alphafold 3 predictions

Integrin α2 and β1 amino acid sequences were obtained from UniProtKB, and propeptides were removed [69,70]. Peptide 1 sequence is provided in **Table 1**. AlphaFold 3 webserver [68] predictions of integrin-peptide interactions, hydroxyprolines specified for Peptide 1. Predictions (25 models across five runs using different seed numbers) identified residues in Peptide in proximity with integrins (≥ 0.5 interaction probability) in at least 20/25 models. Data-analysis was conducted in R v4.4.0. Further model visualization was performed using Mol* Viewer (https://molstar.org/viewer/) version 4.18 (08/06/2025).

### 4.6 Sample collection

Cells were seeded in 6-well plates (50,000 cells/well) and cultured in EGM2 + 10% FBS for 2 days to ∼80% confluency, then starved for 16 hours. Treatments (Peptide 1, COL(I), Peptide 1 + COL(I) or vehicle controls) were applied at migration assay concentrations. Positive controls included 5% FBS and 1mM hydrogen peroxide. Untreated cells received no additions. After 7, 30, or 60 minutes of treatment, plates were placed on ice, washed with ice-cold PBS, and lysed in Laemmli buffer (1X, with 3.6% protease inhibitor, 50 mM sodium fluoride, and 100 μM activated sodium orthovanadate), and cell lysate was collected using cell scrapers. Lysates were bath sonicated for 10 minutes and stored at –20°C. Two biological and two technical replicates were collected (four replicates in total).

### 4.7 Western Blotting

Western blotting was performed as described [98,99] with modifications. Cell extracts (20 μL) were loaded onto polyacrylamide gels until maximum separation was achieved at 40-75 kDa, and proteins transferred to nitrocellulose membranes (Amersham™ Protran^®,^ GE10600004) for 2h on ice. Membranes were blocked in 5% milk-TBS+0.1% Tween™ 20 for 2h, washed with TBS-0.1% Tween™ 20, and incubated overnight at 4°C with primary antibodies ((phospho-ERK, Cell Signaling, Cat# 9106; RRID:AB_331768); ERK, Cell Signaling, Cat# 9107, RRID:AB_10695739; phospho-Akt, Cell Signaling, Cat# 4060, RRID:AB_2315049; 1:1000). Secondary antibodies (1:2000 dilution for both antibodies, Goat Anti-Mouse IgG, HRP; Cell Signaling, Cat# 91196, RRID:AB_2940774, or Goat Anti-Rabbit IgG, HRP; Sigma, Cat# AP132P, RRID:AB_90264; 1:2000) were applied for 2 hours at room temperature. Detection used enhanced chemiluminescence (Perkin and Elmer, NEL104001EA) and Super RX-N films (Fujifilm, 47410 19289). Signals were quantified with Quantity One (Bio-Rad). Phosphoprotein signals were normalized to total protein (ERK) or Coomassie-stained gels (Akt). Analytical gels were run in parallel with their counterparts for western blot, but instead of proceeding to transfer, were incubated with Coomassie brilliant blue overnight and analysed with Quantity One.

### 4.8 LC-MS/MS analysis and MS data processing

Protein extracts (20 μL) were resolved on 12% polyacrylamide gels (5% stacking,) for GeLC-MS/MS [100]. After concentrating the samples in the separating gel using electrophoresis, gels were washed and protein bands were visualized with Coomassie stain, excised, destained (40% acetonitrile, 50 mM ammonium bicarbonate), reduced (10 nM Dithioerythritol in 100 mM ammonium bicarbonate) and alkylated (10 mg/ml iodoacetamide in 100 mM ammonium bicarbonate). Gel pieces were washed with 100 mM ammonium bicarbonate, destain solution and ultrapure water subsequently (each for 20 min at room temperature, while shaking) and dried under vacuum, and trypsinised overnight using 600 ng trypsin per sample in 10 mM ammonium bicarbonate. Peptides were extracted with subsequent incubations in 50 mM ammonium bicarbonate, and twice with 10% formic acid 1:1 with acetonitrile; all three for 15 minutes at room temperature while shaking. Peptide solutions were cleared with a fresh PVDF filter (Merck Milipore) for each sample, and peptides were dried under vacuum. Peptides were reconstituted in mobile phase A (98% water, 2% acetonitrile, 0.1% formic acid) and analysed using a Dionex Ultimate 3000 HPLC nanoflow system coupled to a Q Exactive™ Plus Hybrid Quadrupole-Orbitrap™ mass spectrometer. Samples (10 μL) were loaded onto a Dionex 0.1 x 20 mm, C 18 trap column (flow rate: 5 μL/min) and separated on an Acclaim PepMap^TM^ C18 100 column (75 μm x 50 cm, flow rate 300 nL/min). A 250-minute gradient (2-95% (mobile phase B: 80% acetonitrile, 0.1% formic acid) was applied. Positive ionization was performed at 2.1kV. Data were collected in MS/MS Data Dependent Acquisition mode. Raw files were analysed in Thermo Proteome Discoverer 2.4 using the Sequest search engine and the UniProtKB *Homo sapiens* database (canonical sequences, downloaded 20-06-2019, 20,431 reviewed entries). Carbamidomethylation (cysteine) was a static modification; methionine oxidation and phosphorylation (serine, threonine, tyrosine) were dynamic. Peptide data was filtered (FDR q-value= 0.01; maximum peptide rank = 1). Label-free quantification employed precursor ion peak area. Normalized data were log2-transformed. **Supplementary File**, tab normalized abundance data, provides normalized abundance data. More details on LC-MS/MS and MS data settings are provided in **Appendix A.1.**

### 4.9 Statistical analysis of proteomics results

Phospho-peptides were filtered to retain credible data: peptides with one Peptide-Spectral Match (PSM), lacking unphosphorylated counterparts or with poor spectra were excluded (**Supplementary File**, tab removed peptides**, Supplementary** Figure 4). Spectra were manually inspected based on defined criteria [83,84]. Retained peptides fulfilled ≥3/4 criteria. Details are in **Supplementary File**, tab spectra analysis. Peptide and COL(I)-treated cells (n=4 /time point) were compared to COL(I)-only treated cells. Time-independent comparisons applied Benjamini-Hochberg (BH) correction to p-values. Wilcoxon rank sum tests evaluated differential abundance. Analyses were performed in R v4.4.0. More details on peptide filtering are provided in **Appendix A.2**.

## 5. Conclusions

Our results demonstrate that a naturally occurring COLα1 (I)-derived peptide can inhibit collagen-induced endothelial migration at a physiological concentration, probably by interacting with the integrin α2β1 receptor. It should be noted that the interaction of peptide 1 with this receptor, while supported by the AlphaFold3 predictions and phosphoproteomics results, has not yet been experimentally confirmed. In addition, key signalling molecules involved in migration are downregulated in COLα1 (I) + Peptide 1 treated cells, when compared to COLα1 (I) alone, further supporting the classification of Peptide 1 as a matrikine. However, further investigation to elucidate the exact underlying signalling mechanism is needed.

These findings open new avenues for research into the detailed characterization of Peptide 1 mechanism of action and its therapeutic potential. Future studies should also explore the mechanisms underlying its urinary excretion, which may be linked to its biological activity, which would aid in understanding the peptide’s role as a biomarker in diseases associated with endothelial dysfunction.

## Supplementary Materials

The following supporting information can be downloaded at: https://www.mdpi.com/article/doi/s1: Supplementary File 1 (proteomics results and unprocessed data) and Supplementary Figures (migration results, western blots).

## Author Contributions

Conceptualization, Ioanna K. Mina, Marika Mokou, Jerome Zoidakis, Harald Mischak, Antonia Vlahou and Agnieszka Latosinska; Data curation, Hanne Devos; Formal analysis, Hanne Devos, Jerome Zoidakis and Maria G. Roubelakis; Funding acquisition, Harald Mischak, Antonia Vlahou and Agnieszka Latosinska; Investigation, Hanne Devos, Ioanna K. Mina, Foteini Paradeisi, Manousos Makridakis, Aggeliki Tserga, Agnieszka Latosinska and Maria G. Roubelakis; Methodology, Ioanna K. Mina, Foteini Paradeisi, Manousos Makridakis, Aggeliki Tserga, Marika Mokou, Jerome Zoidakis, Harald Mischak, Antonia Vlahou, Agnieszka Latosinska and Maria G. Roubelakis; Project administration, Antonia Vlahou and Agnieszka Latosinska; Resources, Antonia Vlahou and Maria G. Roubelakis; Supervision, Jerome Zoidakis, Harald Mischak, Antonia Vlahou, Agnieszka Latosinska and Maria G. Roubelakis; Validation, Hanne Devos, Ioanna K. Mina and Antonia Vlahou; Visualization, Hanne Devos; Writing – original draft, Hanne Devos; Writing – review & editing, Hanne Devos, Marika Mokou, Jerome Zoidakis, Harald Mischak, Antonia Vlahou, Agnieszka Latosinska and Maria G. Roubelakis.

## Institutional Review Board Statement

Not applicable

## Informed Consent Statement

Not applicable

## Data Availability Statement

Raw proteomics data files have been deposited on MassIVE (identifier: MSV000097296; username for reviewer access: MSV000097296_reviewer; password: dV5qsfE4rYv37q81) as well as ProteomExchange (PXD061753) and Supplementary File 1. A Proteom Exchange dataset identifier has been requested. The experimental metadata file has been generated using lesSDRF [101]. Supplementary File 1 is similarly available through MassIVE under the same identifier.

## Acknowledgments

**Figure 1** and **figure 5** were made with the aid of Servier Medical Art (CC BY 4.0 license).

## Abbreviations

The following abbreviations are used in this manuscript:

7-AAD: 7-Aminoactinomycin D
BSA: Bovine Serum Albumin
CE-MS/MS: capillary electrophoresis combined with mass spectrometry
CKD: Chronic kidney disease
COL(I): Collagen type I
COLα1 (I): Collagen type I, alpha 1 chain
COLα2 (I): Collagen type I, alpha 2 chain
COPD: chronic obstructive pulmonary disease
CXCL: C-X-C Ligand
CXCR: C-X-C Chemokine receptor
EBM2: Endothelial Basal Medium
ECM: Extracellular Matrix
EDTA: Ethylenediaminetetraacetic acid
EGM2: Endothelial Growth Medium
ERK: Extracellular Regulated Kinase
FAK: Focal Adhesion Kinase
FBS: Fetal Bovine Serum
FDR: False Discovery Rate
FITC: Fluorescein isothiocyanate
HUVEC: Human Umbilical Vein Endothelial Cells
IP3: 1,4,5-triphosphate
JNK: C-jun N-terminal kinase
LC-MS/MS: Liquid Chromatography – Mass spectrometry
MMP: Matrix-metalloproteinases
PAK: P21 (Rac1) Activated Kinase
PE: phycoerythrin
PFA: Paraformaldehyde
PBS: Phosphate Buffered Saline
PKC: Protein Kinase C
PLC: Phospholipase C
PRAK: P38-regulated/activated protein kinase PSM Peptide-spectral match
STAT: Signal Transducer and Activator of Transcription
TBS: Tris-buffered Saline
VEGF: Vascular Endothelial Growth Factor

## Appendix A

### Appendix A.1

For LC-MS/MS analysis a Dionex Ultimate 3000 HPLC nanoflow system was used in combination with a Q Exactive™ Plus Hybrid Quadrupole-Orbitrap™ mass spectrometer. Samples were reconstituted in 25μL mobile phase A (98% water, 2% acetonitrile, 0,1% formic acid) and 10μL was loaded onto the LC column configured with a Dionex 0.1 x 20 mm, 5 μm, 100 Ä C 18 nano trap column. A flow rate of 5 μL/min was used. An Acclaim PepMap^TM^ C18 100 nanoViper column 75 μm x 50 cm, 2 μm 100 Ä with a flow rate of 300nL/min was used as analytical column. The column was washed and re-equilibrated prior to each sample injection. Mobile phase B consisted of 0.1% formic acid in 80% acetonitrile. The eluent was ionized using a nanospray Flex^TM^ ESI source in positive ion mode, whereas the Q Exactive Orbitrap (Thermo Finnigan, Bremen, Germany), was operated in MS/MS mode. Peptides were eluted under a 250min gradient from 2% mobile phase B to 95% mobile phase B. Positive ion electrospray ionization with a voltage of 2.1 kV was used for gaseous phase transition of eluted, separated peptides. The mass spectrometer parameters were set as follows: 1) full MS: resolution of 70,000, automatic gain control (AGC) target at 1e6, maximum injection time (IT) of 100 ms, scan range of 380 – 1,200 m/z; 2) dd-MS^2^: resolution of 17.500, AGC target at 1e5, maximum IT of 50ms, TopN of 20, normalized collision energy of 30; 3) dd settings: intensity threshold 1.6e5, Minimum AGC target 8.00e3, dynamic exclusion for 15s.

Raw files were processed with Thermo Proteome Discoverer 2.4 software, utilizing the Sequest search engine and the UniProt *Homo sapiens* fasta database containing only canonical sequences (downloaded on 20-06-2019, including 20,431 reviewed entries). Carbamidomethylation of cysteine was set as a static modification, and oxidation of methionine and phosphorylation of serine, threonine and tyrosine were set as dynamic modifications. Two missed cleavage sites, a precursor mass tolerance of 5 ppm, and fragment mass tolerance of 0.05 Da were allowed. Feature mapper node was run with default settings. The following filters were also used: peptide: High Confidence (FDR q value based = 0.01), peptide rank: Maximum rank = 1, peptide grouping: enabled, protein grouping: enabled. For quantification, the label-free method was employed using the peak area of precursor ions to determine the relative abundance of identified proteins. the label-free method was employed using the peak area of precursor ions to determine the relative abundance of identified proteins and peptides. Missing values were replaced by zeros. Part per million (ppm) normalization was applied once at the protein level to evaluate the presence of the integrin receptor and once at the peptide level to evaluate phosphorylation events, by dividing the peptide summed peak area by the total peak area in a sample, multiplied by one million. Normalized abundance data was then log2-transformed, and, for phospho-peptides, can be found in Supplementary File 1, tab “normalized abundance data”.

### Appendix A.2

Phosphorylated peptides’ credibility was assessed as follows: first, all peptides with only one Peptide-Spectral Match (PSM) were discarded (n=21), peptides without unphosphorylated peptide mapping to the same protein, were removed (n=6), and peptides with poor spectra (n=7) were filtered out (**Supplementary File 1**, tab “removed peptides”; mass spectra can be found in **Supplementary** Figure 2). The criteria for the manual inspection of the spectra were: a) four sequential, explained fragments had to be observed or five explained fragments, separated by maximum one unexplained fragment, in a series of six sequential fragments had to be observed; b) the b-or y-ion corresponding to the breakage of the proline peptide bond had to have an intensity of at least 50% of the most intense ion observed in the spectrum; c) at least six out of the ten most intense ions had to correspond to b-or y-ions; d) at least three ions corresponding to the neutral loss of the phospho-group had to be observed. For the last rule, an exception was made if the peptide contained multiple prolines, as this has been shown to affect the neutral loss of the phospho-group [84]. Peptides of which the spectra fulfil at least three out of four criteria, based on the work of Potel et al. [83], were retained.

## Appendix B

**Table B1:**
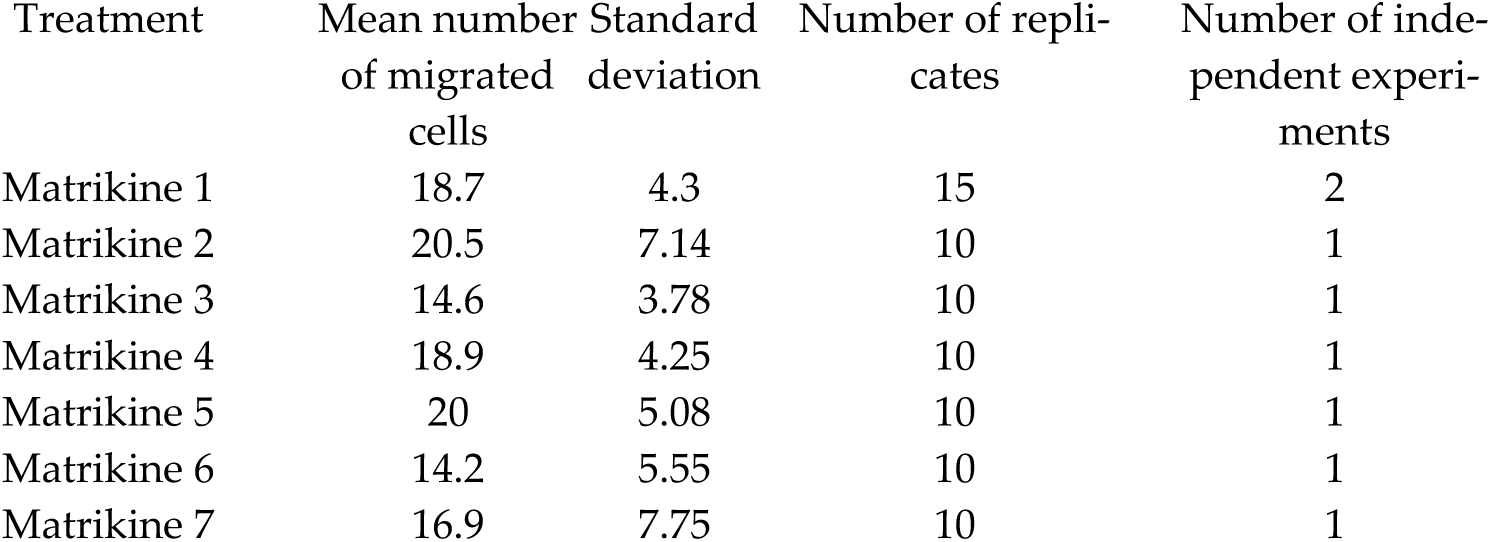
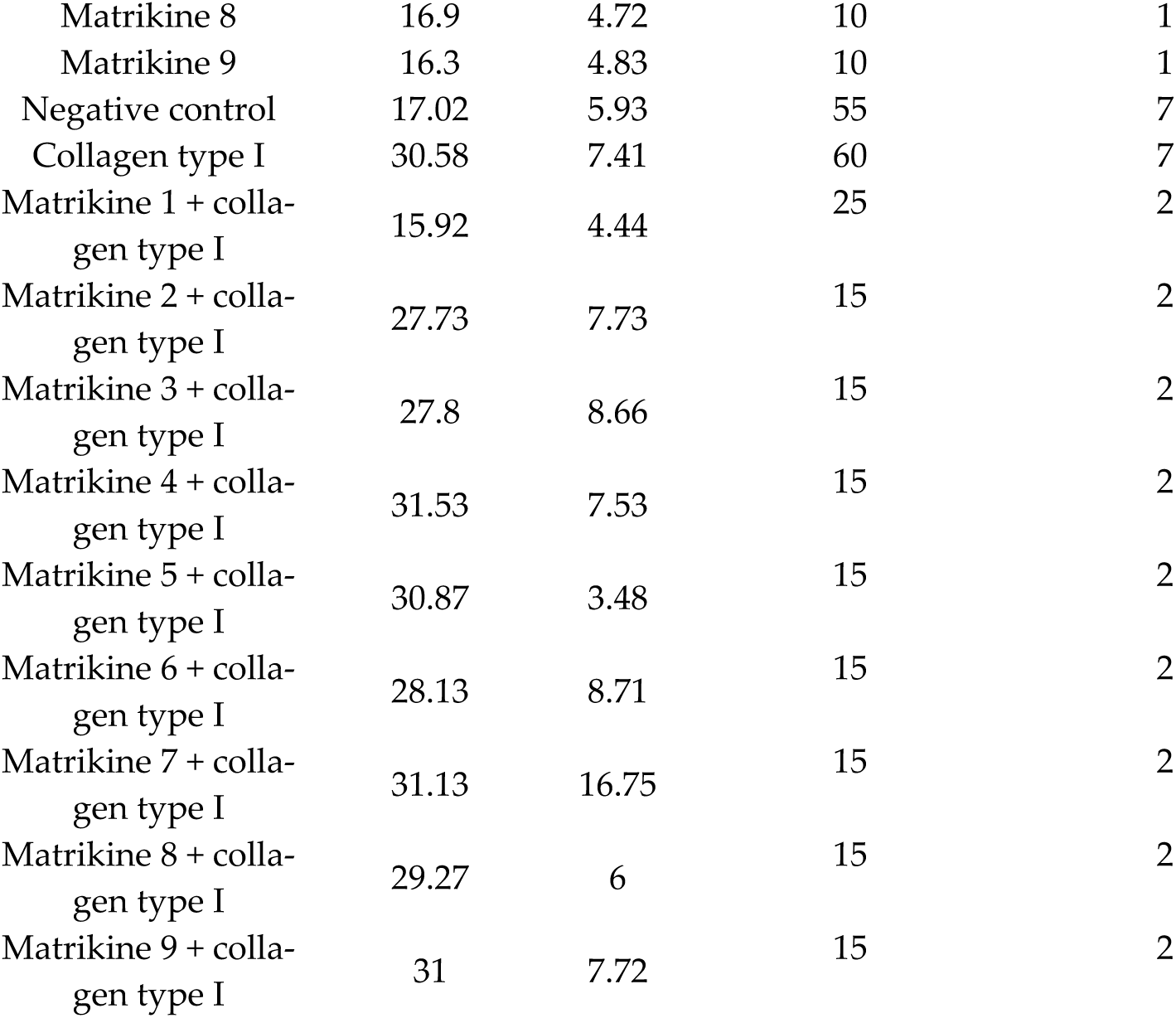
Overview of the mean number of migrated cells and their standard deviation, for each of the peptides, their combination with full-length COL(I), COL(I) by itself and negative control. The number of replicates (biological and technical) is additionally reported. Lastly, the number of independent experiment (experiments conducted on different days, using cells of the same passage number) has additionally been provided.

| Treatment | Mean number of migrated cells | Standard deviation | Number of replicates | Number of independent experiments |
| --- | --- | --- | --- | --- |
| Matrikine 1 | 18.7 | 4.3 | 15 | 2 |
| Matrikine 2 | 20.5 | 7.14 | 10 | 1 |
| Matrikine 3 | 14.6 | 3.78 | 10 | 1 |
| Matrikine 4 | 18.9 | 4.25 | 10 | 1 |
| Matrikine 5 | 20 | 5.08 | 10 | 1 |
| Matrikine 6 | 14.2 | 5.55 | 10 | 1 |
| Matrikine 7 | 16.9 | 7.75 | 10 | 1 |
| Matrikine 8 | 16.9 | 4.72 | 10 | 1 |
| Matrikine 9 | 16.3 | 4.83 | 10 | 1 |
| Negative control | 17.02 | 5.93 | 55 | 7 |
| Collagen type I | 30.58 | 7.41 | 60 | 7 |
| Matrikine 1 + collagen type I | 15.92 | 4.44 | 25 | 2 |
| Matrikine 2 + collagen type I | 27.73 | 7.73 | 15 | 2 |
| Matrikine 3 + collagen type I | 27.8 | 8.66 | 15 | 2 |
| Matrikine 4 + collagen type I | 31.53 | 7.53 | 15 | 2 |
| Matrikine 5 + collagen type I | 30.87 | 3.48 | 15 | 2 |
| Matrikine 6 + collagen type I | 28.13 | 8.71 | 15 | 2 |
| Matrikine 7 + collagen type I | 31.13 | 16.75 | 15 | 2 |
| Matrikine 8 + collagen type I | 29.27 | 6 | 15 | 2 |
| Matrikine 9 + collagen type I | 31 | 7.72 | 15 | 2 |

## Disclaimer/Publisher’s Note

The statements, opinions and data contained in all publications are solely those of the individual author(s) and contributor(s) and not of MDPI and/or the editor(s). MDPI and/or the editor(s) disclaim responsibility for any injury to people or property resulting from any ideas, methods, instructions or products referred to in the content.

## References

1. Theocharis, A.D.; Manou, D.; Karamanos, N.K. The Extracellular Matrix as a Multitasking Player in Disease. FEBS J. 2019, 286, 2830–2869, doi:10.1111/febs.14818.

2. Devos, H.; Zoidakis, J.; Roubelakis, M.G.; Latosinska, A.; Vlahou, A. Reviewing the Regulators of COL1A1. Int. J. Mol. Sci. 2023, 24, 10004, doi:10.3390/ijms241210004.

3. Shi, R.; Zhang, Z.; Zhu, A.; Xiong, X.; Zhang, J.; Xu, J.; Sy, M.-S.; Li, C. Targeting Type I Collagen for Cancer Treatment. Int. J. Cancer 2022, 151, 665–683, doi:10.1002/ijc.33985.

4. Di Lullo, G.A.; Sweeney, S.M.; Korkko, J.; Ala-Kokko, L.; San Antonio, J.D. Mapping the Ligand-Binding Sites and Disease-Associated Mutations on the Most Abundant Protein in the Human, Type I Collagen. J. Biol. Chem. 2002, 277, 4223–4231, doi:10.1074/jbc.M110709200.

5. Li, S.; Sampson, C.; Liu, C.; Piao, H.; Liu, H.-X. Integrin Signaling in Cancer: Bidirectional Mechanisms and Therapeutic Opportunities. Cell Commun. Signal. 2023, 21, 266, doi:10.1186/s12964-023-01264-4.

6. Sun, L.; Guo, S.; Xie, Y.; Yao, Y. The Characteristics and the Multiple Functions of Integrin Β1 in Human Cancers. J. Transl. Med. 2023, 21, 787, doi:10.1186/s12967-023-04696-1.

7. Aman, J.; Margadant, C. Integrin-Dependent Cell–Matrix Adhesion in Endothelial Health and Disease. Circ. Res. 2023, 132, 355–378, doi:10.1161/CIRCRESAHA.122.322332.

8. Godoy, P.; Hengstler, J.G.; Ilkavets, I.; Meyer, C.; Bachmann, A.; Müller, A.; Tuschl, G.; Mueller, S.O.; Dooley, S. Extracellular Matrix Modulates Sensitivity of Hepatocytes to Fibroblastoid Dedifferentiation and Transforming Growth Factor Beta-Induced Apoptosis. Hepatol. Baltim. Md 2009, 49, 2031–2043, doi:10.1002/hep.22880.

9. Viale-Bouroncle, S.; Gosau, M.; Morsczeck, C. Collagen I Induces the Expression of Alkaline Phosphatase and Osteopontin via Independent Activations of FAK and ERK Signalling Pathways. Arch. Oral Biol. 2014, 59, 1249– 1255, doi:10.1016/j.archoralbio.2014.07.013.

10. Krasny, L.; Shimony, N.; Tzukert, K.; Gorodetsky, R.; Lecht, S.; Nettelbeck, D.M.; Haviv, Y.S. An In-Vitro Tumour Microenvironment Model Using Adhesion to Type I Collagen Reveals Akt-Dependent Radiation Resistance in Renal Cancer Cells. Nephrol. Dial. Transplant. 2010, 25, 373–380, doi:10.1093/ndt/gfp525.

11. Jarvis, G.E.; Best, D.; Watson, S.P. Glycoprotein VI/Fc Receptor γ Chain-Independent Tyrosine Phosphorylation and Activation of Murine Platelets by Collagen. Biochem. J. 2004, 383, 581–588, doi:10.1042/BJ20040654.

12. Takeuchi, Y.; Suzawa, M.; Kikuchi, T.; Nishida, E.; Fujita, T.; Matsumoto, T. Differentiation and Transforming Growth Factor-β Receptor Down-Regulation by Collagen-Α2β1 Integrin Interaction Is Mediated by Focal Adhesion Kinase and Its Downstream Signals in Murine Osteoblastic Cells *. J. Biol. Chem. 1997, 272, 29309–29316, doi:10.1074/jbc.272.46.29309.

13. Ziaee, S.; Chung, L.W. Induction of Integrin Α2 in a Highly Bone Metastatic Human Prostate Cancer Cell Line: Roles of RANKL and AR under Three-Dimensional Suspension Culture. Mol. Cancer 2014, 13, 208, doi:10.1186/1476-4598-13-208.

14. Stejskalová, A.; Fincke, V.; Nowak, M.; Schmidt, Y.; Borrmann, K.; von Wahlde, M.-K.; Schäfer, S.D.; Kiesel, L.; Greve, B.; Götte, M. Collagen I Triggers Directional Migration, Invasion and Matrix Remodeling of Stroma Cells in a 3D Spheroid Model of Endometriosis. Sci. Rep. 2021, 11, 4115, doi:10.1038/s41598-021-83645-8.

15. Goetsch, K.P.; Kallmeyer, K.; Niesler, C.U. Decorin Modulates Collagen I-Stimulated, but Not Fibronectin-Stimulated, Migration of C2C12 Myoblasts. Matrix Biol. 2011, 30, 109–117, doi:10.1016/j.matbio.2010.10.009.

16. Jabłońska-Trypuć, A.; Matejczyk, M.; Rosochacki, S. Matrix Metalloproteinases (MMPs), the Main Extracellular Matrix (ECM) Enzymes in Collagen Degradation, as a Target for Anticancer Drugs. J. Enzyme Inhib. Med. Chem. 2016, 31, 177–183, doi:10.3109/14756366.2016.1161620.

17. Vizovišek, M.; Fonović, M.; Turk, B. Cysteine Cathepsins in Extracellular Matrix Remodeling: Extracellular Matrix Degradation and Beyond. Matrix Biol. J. Int. Soc. Matrix Biol. 2019, 75–76, 141–159, doi:10.1016/j.matbio.2018.01.024.

18. Jariwala, N.; Ozols, M.; Bell, M.; Bradley, E.; Gilmore, A.; Debelle, L.; Sherratt, M.J. Matrikines as Mediators of Tissue Remodelling. Adv. Drug Deliv. Rev. 2022, 185, 114240, doi:10.1016/j.addr.2022.114240.

19. Latosinska, A.; Siwy, J.; Faguer, S.; Beige, J.; Mischak, H.; Schanstra, J.P. Value of Urine Peptides in Assessing Kidney and Cardiovascular Disease. PROTEOMICS – Clin. Appl. 2021, 15, 2000027, doi:10.1002/prca.202000027.

20. Latosinska, A.; Siwy, J.; Mischak, H.; Frantzi, M. Peptidomics and Proteomics Based on CE-MS as a Robust Tool in Clinical Application: The Past, the Present, and the Future. Electrophoresis 2019, 40, 2294–2308, doi:10.1002/elps.201900091.

21. Staatz, W.D.; Fok, K.F.; Zutter, M.M.; Adams, S.P.; Rodriguez, B.A.; Santoro, S.A. Identification of a Tetrapeptide Recognition Sequence for the Alpha 2 Beta 1 Integrin in Collagen. J. Biol. Chem. 1991, 266, 7363–7367, doi:10.1016/S0021-9258(20)89455-1.

22. Cha, B.-H.; Shin, S.R.; Leijten, J.; Li, Y.-C.; Singh, S.; Liu, J.C.; Annabi, N.; Abdi, R.; Dokmeci, M.R.; Vrana, N.E.;, et al. Integrin-Mediated Interactions Control Macrophage Polarization in 3D Hydrogels. Adv. Healthc. Mater. 2017, *6*, 10.1002/adhm.201700289, doi:10.1002/adhm.201700289.

23. Mehta, M.; Madl, C.M.; Lee, S.; Duda, G.N.; Mooney, D.J. The Collagen I Mimetic Peptide DGEA Enhances an Osteogenic Phenotype in Mesenchymal Stem Cells When Presented from Cell-Encapsulating Hydrogels. J. Biomed. Mater. Res. A 2015, 103, 3516–3525, doi:10.1002/jbm.a.35497.

24. Madamanchi, A.; Santoro, S.A.; Zutter, M.M. Α2β1 Integrin. Adv. Exp. Med. Biol. 2014, 819, 41–60, doi:10.1007/978-94-017-9153-3_3.

25. Mineur, P.; Guignandon, A.; Lambert, Ch.A.; Amblard, M.; Lapière, Ch.M.; Nusgens, B.V. RGDS and DGEA-Induced [Ca2+]i Signalling in Human Dermal Fibroblasts. Biochim. Biophys. Acta BBA – Mol. Cell Res. 2005, 1746, 28–37, doi:10.1016/j.bbamcr.2005.07.004.

26. McCann, T.J.; Terranova, G.; Keyte, J.W.; Papaioannou, S.S.; Mason, W.T.; Meikle, M.C.; McDonald, F. An Analysis of Ca2+ Release by DGEA: Mobilization of Two Functionally Distinct Internal Stores in Saos-2 Cells. Am. J. Physiol.-Cell Physiol. 1998, 275, C33–C41, doi:10.1152/ajpcell.1998.275.1.C33.

27. McCann, T.J.; Mason, W.T.; Meikle, M.C.; McDonald, F. A Collagen Peptide Motif Activates Tyrosine Kinase-Dependent Calcium Signalling Pathways in Human Osteoblast-like Cells. Matrix Biol. 1997, 16, 273–283, doi:10.1016/S0945-053X(97)90015-9.

28. Kim, H.-K.; Joe, Y.A. DGDA, a Local Sequence of the Kringle 2 Domain, Is a Functional Motif of the Tissue-Type Plasminogen Activator’s Antiangiogenic Kringle Domain. Biochem. Biophys. Res. Commun. 2010, 391, 166–169, doi:10.1016/j.bbrc.2009.11.025.

29. Chen, S.; Chakrabarti, R.; Keats, E.C.; Chen, M.; Chakrabarti, S.; Khan, Z.A. Regulation of Vascular Endothelial Growth Factor Expression by Extra Domain B Segment of Fibronectin in Endothelial Cells. Invest. Ophthalmol. Vis. Sci. 2012, 53, 8333–8343, doi:10.1167/iovs.12-9766.

30. Jha, A.; Moore, E. Collagen-Derived Peptide, DGEA, Inhibits pro-Inflammatory Macrophages in Biofunctional Hydrogels. J. Mater. Res. 2022, 37, 77–87, doi:10.1557/s43578-021-00423-y.

31. Anderson, J.M.; Vines, J.B.; Patterson, J.L.; Chen, H.; Javed, A.; Jun, H.-W. Osteogenic Differentiation of Human Mesenchymal Stem Cells Synergistically Enhanced by Biomimetic Peptide Amphiphiles Combined with Conditioned Medium. Acta Biomater. 2011, 7, 675–682, doi:10.1016/j.actbio.2010.08.016.

32. Chen, G.; Kong, P.; Jiang, A.; Wang, X.; Sun, Y.; Yu, T.; Chi, H.; Song, C.; Zhang, H.; Subedi, D.;, et al. A Modular Programmed Biphasic Dual-Delivery System on 3D Ceramic Scaffolds for Osteogenesis in Vitro and in Vivo. J. Mater. Chem. B 2020, 8, 9697–9717, doi:10.1039/C9TB02127B.

33. Hamaia, S.; Farndale, R.W. Integrin Recognition Motifs in the Human Collagens. Adv. Exp. Med. Biol. 2014, 819, 127–142, doi:10.1007/978-94-017-9153-3_9.

34. Knight, C.G.; Morton, L.F.; Peachey, A.R.; Tuckwell, D.S.; Farndale, R.W.; Barnes, M.J. The Collagen-Binding A-Domains of Integrins Α1β1 and α2β1Recognize the Same Specific Amino Acid Sequence, GFOGER, in Native (Triple-Helical) Collagens *. J. Biol. Chem. 2000, 275, 35–40, doi:10.1074/jbc.275.1.35.

35. Reyes, C.D.; García, A.J. Α2β1 Integrin-Specific Collagen-Mimetic Surfaces Supporting Osteoblastic Differentiation. J. Biomed. Mater. Res. A 2004, 69A, 591–600, doi:10.1002/jbm.a.30034.

36. Fraser, D.; Benoit, D. Dual Peptide-Functionalized Hydrogels Differentially Control Periodontal Cell Function and Promote Tissue Regeneration. Biomater. Adv. 2022, 141, 213093, doi:10.1016/j.bioadv.2022.213093.

37. Clark, A.Y.; Martin, K.E.; García, J.R.; Johnson, C.T.; Theriault, H.S.; Han, W.M.; Zhou, D.W.; Botchwey, E.A.; García, A.J. Integrin-Specific Hydrogels Modulate Transplanted Human Bone Marrow-Derived Mesenchymal Stem Cell Survival, Engraftment, and Reparative Activities. Nat. Commun. 2020, 11, 114, doi:10.1038/s41467-019-14000-9.

38. Consonni, A.; Cipolla, L.; Guidetti, G.; Canobbio, I.; Ciraolo, E.; Hirsch, E.; Falasca, M.; Okigaki, M.; Balduini, C.; Torti, M. Role and Regulation of Phosphatidylinositol 3-Kinase β in Platelet Integrin Α2β1 Signaling. Blood 2012, 119, 847–856, doi:10.1182/blood-2011-07-364992.

39. Sweeney, S.M.; DiLullo, G.; Slater, S.J.; Martinez, J.; Iozzo, R.V.; Lauer-Fields, J.L.; Fields, G.B.; Antonio, J.D.S. Angiogenesis in Collagen I Requires Α2β1 Ligation of a GFP*GER Sequence and Possibly P38 MAPK Activation and Focal Adhesion Disassembly *. J. Biol. Chem. 2003, 278, 30516–30524, doi:10.1074/jbc.M304237200.

40. Taubenberger, A.V.; Bray, L.J.; Haller, B.; Shaposhnykov, A.; Binner, M.; Freudenberg, U.; Guck, J.; Werner, C. 3D Extracellular Matrix Interactions Modulate Tumour Cell Growth, Invasion and Angiogenesis in Engineered Tumour Microenvironments. Acta Biomater. 2016, 36, 73–85, doi:10.1016/j.actbio.2016.03.017.

41. Patel, D.F.; Snelgrove, R.J. The Multifaceted Roles of the Matrikine Pro-Gly-Pro in Pulmonary Health and Disease. Eur. Respir. Rev. 2018, 27, 180017, doi:10.1183/16000617.0017-2018.

42. Misiura, M.; Miltyk, W. Proline-Containing Peptides-New Insight and Implications: A Review. BioFactors Oxf. Engl. 2019, 45, 857–866, doi:10.1002/biof.1554.

43. Weathington, N.M.; van Houwelingen, A.H.; Noerager, B.D.; Jackson, P.L.; Kraneveld, A.D.; Galin, F.S.; Folkerts, G.; Nijkamp, F.P.; Blalock, J.E. A Novel Peptide CXCR Ligand Derived from Extracellular Matrix Degradation during Airway Inflammation. Nat. Med. 2006, 12, 317–323, doi:10.1038/nm1361.

44. Hahn, C.S.; Scott, D.W.; Xu, X.; Roda, M.A.; Payne, G.A.; Wells, J.M.; Viera, L.; Winstead, C.J.; Bratcher, P.; Sparidans, R.W.;, et al. The Matrikine N-α-PGP Couples Extracellular Matrix Fragmentation to Endothelial Permeability. Sci. Adv. 2015, 1, e1500175, doi:10.1126/sciadv.1500175.

45. Li, X.; Tang, Y.; Yu, F.; Sun, Y.; Huang, F.; Chen, Y.; Yang, Z.; Ding, G. Inhibition of Prostate Cancer DU-145 Cells Proliferation by Anthopleura Anjunae Oligopeptide (YVPGP) via PI3K/AKT/mTOR Signaling Pathway. Mar. Drugs 2018, 16, 325, doi:10.3390/md16090325.

46. Feng, C.; Zhang, Y.; Yang, M.; Huang, B.; Zhou, Y. Collagen-Derived N-Acetylated Proline-Glycine-Proline in Intervertebral Discs Modulates CXCR1/2 Expression and Activation in Cartilage Endplate Stem Cells to Induce Migration and Differentiation Toward a Pro-Inflammatory Phenotype. Stem Cells Dayt. Ohio 2015, 33, 3558–3568, doi:10.1002/stem.2200.

47. Kwon, Y.W.; Heo, S.C.; Lee, T.W.; Park, G.T.; Yoon, J.W.; Jang, I.H.; Kim, S.-C.; Ko, H.-C.; Ryu, Y.; Kang, H.;, et al. N-Acetylated Proline-Glycine-Proline Accelerates Cutaneous Wound Healing and Neovascularization by Human Endothelial Progenitor Cells. Sci. Rep. 2017, 7, 43057, doi:10.1038/srep43057.

48. Thaler, R.; Zwerina, J.; Rumpler, M.; Spitzer, S.; Gamsjaeger, S.; Paschalis, E.P.; Klaushofer, K.; Varga, F. Homocysteine Induces Serum Amyloid A3 in Osteoblasts via Unlocking RGD-Motifs in Collagen. FASEB J. 2013, 27, 446–463, doi:10.1096/fj.12-208058.

49. Taubenberger, A.V.; Woodruff, M.A.; Bai, H.; Muller, D.J.; Hutmacher, D.W. The Effect of Unlocking RGD-Motifs in Collagen I on Pre-Osteoblast Adhesion and Differentiation. Biomaterials 2010, 31, 2827–2835, doi:10.1016/j.biomaterials.2009.12.051.

50. Davis, G.E. Affinity of Integrins for Damaged Extracellular Matrix: Αvβ3 Binds to Denatured Collagen Type I through RGD Sites. Biochem. Biophys. Res. Commun. 1992, 182, 1025–1031, doi:10.1016/0006-291X(92)91834-D.

51. Davis, G.E.; Bayless, K.J.; Davis, M.J.; Meininger, G.A. Regulation of Tissue Injury Responses by the Exposure of Matricryptic Sites within Extracellular Matrix Molecules. Am. J. Pathol. 2000, 156, 1489–1498, doi:10.1016/S0002-9440(10)65020-1.

52. Ma, Z.; Mao, C.; Jia, Y.; Fu, Y.; Kong, W. Extracellular Matrix Dynamics in Vascular Remodeling. Am. J. Physiol. – Cell Physiol. 2020, 319, C481–C499, doi:10.1152/ajpcell.00147.2020.

53. Argilés, Á.; Siwy, J.; Duranton, F.; Gayrard, N.; Dakna, M.; Lundin, U.; Osaba, L.; Delles, C.; Mourad, G.; Weinberger, K.M.;, et al. CKD273, a New Proteomics Classifier Assessing CKD and Its Prognosis. PloS One 2013, 8, e62837, doi:10.1371/journal.pone.0062837.

54. Hobson, S.; Mavrogeorgis, E.; He, T.; Siwy, J.; Ebert, T.; Kublickiene, K.; Stenvinkel, P.; Mischak, H. Urine Peptidome Analysis Identifies Common and Stage-Specific Markers in Early Versus Advanced CKD. Proteomes 2023, 11, 25, doi:10.3390/proteomes11030025.

55. Campbell, R.T.; Jasilek, A.; Mischak, H.; Nkuipou-Kenfack, E.; Latosinska, A.; Welsh, P.I.; Jackson, C.E.; Cannon, J.; McConnachie, A.; Delles, C.;, et al. The Novel Urinary Proteomic Classifier HF1 Has Similar Diagnostic and Prognostic Utility to BNP in Heart Failure. ESC Heart Fail. 2020, 7, 1595–1604, doi:10.1002/ehf2.12708.

56. Bannaga, A.S.; Metzger, J.; Kyrou, I.; Voigtländer, T.; Book, T.; Melgarejo, J.; Latosinska, A.; Pejchinovski, M.; Staessen, J.A.; Mischak, H.;, et al. Discovery, Validation and Sequencing of Urinary Peptides for Diagnosis of Liver Fibrosis-A Multicentre Study. EBioMedicine 2020, 62, 103083, doi:10.1016/j.ebiom.2020.103083.

57. Diaz-Ricart, M.; Torramade-Moix, S.; Pascual, G.; Palomo, M.; Moreno-Castaño, A.B.; Martinez-Sanchez, J.; Vera, M.; Cases, A.; Escolar, G. Endothelial Damage, Inflammation and Immunity in Chronic Kidney Disease. Toxins 2020, 12, 361, doi:10.3390/toxins12060361.

58. Godo, S.; Shimokawa, H. Endothelial Functions. Arterioscler. Thromb. Vasc. Biol. 2017, 37, e108–e114, doi:10.1161/ATVBAHA.117.309813.

59. Magalhães, P.; Pontillo, C.; Pejchinovski, M.; Siwy, J.; Krochmal, M.; Makridakis, M.; Carrick, E.; Klein, J.; Mullen, W.; Jankowski, J.;, et al. Comparison of Urine and Plasma Peptidome Indicates Selectivity in Renal Peptide Handling. Proteomics Clin. Appl. 2018, 12, e1700163, doi:10.1002/prca.201700163.

60. Ganapathy, V.; Leibach, F.H. Carrier-Mediated Reabsorption of Small Peptides in Renal Proximal Tubule. Am. J. Physiol.-Ren. Physiol. 1986, 251, F945–F953, doi:10.1152/ajprenal.1986.251.6.F945.

61. Fiedler, L.R.; Schönherr, E.; Waddington, R.; Niland, S.; Seidler, D.G.; Aeschlimann, D.; Eble, J.A. Decorin Regulates Endothelial Cell Motility on Collagen I through Activation of Insulin-like Growth Factor I Receptor and Modulation of Alpha2beta1 Integrin Activity. J. Biol. Chem. 2008, 283, 17406–17415, doi:10.1074/jbc.M710025200.

62. Nakamura, J.; Shigematsu, S.; Yamauchi, K.; Takeda, T.; Yamazaki, M.; Kakizawa, T.; Hashizume, K. Biphasic Function of Focal Adhesion Kinase in Endothelial Tube Formation Induced by Fibril-Forming Collagens. Biochem. Biophys. Res. Commun. 2008, 374, 699–703, doi:10.1016/j.bbrc.2008.07.123.

63. Senger, D.R.; Perruzzi, C.A.; Streit, M.; Koteliansky, V.E.; de Fougerolles, A.R.; Detmar, M. The Α1β1 and Α2β1 Integrins Provide Critical Support for Vascular Endothelial Growth Factor Signaling, Endothelial Cell Migration, and Tumor Angiogenesis. Am. J. Pathol. 2002, 160, 195–204, doi:10.1016/S0002-9440(10)64363-5.

64. de Kruijf, P.; Lim, H.D.; Overbeek, S.A.; Zaman, G.J.R.; Kraneveld, A.D.; Folkerts, G.; Leurs, R.; Smit, M.J. The Collagen-Breakdown Product *N*-Acetyl-Proline-Glycine-Proline (*N-*α*-*PGP) Does Not Interact Directly with Human CXCR1 and CXCR2. Eur. J. Pharmacol. 2010, 643, 29–33, doi:10.1016/j.ejphar.2010.06.017.

65. Knight, C.G.; Morton, L.F.; Onley, D.J.; Peachey, A.R.; Messent, A.J.; Smethurst, P.A.; Tuckwell, D.S.; Farndale, R.W.; Barnes, M.J. Identification in Collagen Type I of an Integrin Α2β1-Binding Site Containing an Essential GER Sequence *. J. Biol. Chem. 1998, 273, 33287–33294, doi:10.1074/jbc.273.50.33287.

66. Siwy, J.; Mullen, W.; Golovko, I.; Franke, J.; Zürbig, P. Human Urinary Peptide Database for Multiple Disease Biomarker Discovery. PROTEOMICS – Clin. Appl. 2011, 5, 367–374, doi:10.1002/prca.201000155.

67. COL1A1-Collagen Alpha-1(I) Chain – Homo Sapiens (Human) | UniProtKB | UniProt Available online: https://www.uniprot.org/uniprotkb/P02452/entry (accessed on 9 January 2023).

68. Abramson, J.; Adler, J.; Dunger, J.; Evans, R.; Green, T.; Pritzel, A.; Ronneberger, O.; Willmore, L.; Ballard, A.J.; Bambrick, J.;, et al. Accurate Structure Prediction of Biomolecular Interactions with AlphaFold 3. Nature 2024, 630, 493–500, doi:10.1038/s41586-024-07487-w.

69. ITGA2 – Integrin Alpha-2 – Homo Sapiens (Human) | UniProtKB | UniProt Available online: https://www.uniprot.org/uniprotkb/P17301/entry (accessed on 19 September 2024).

70. ITGB1 – Integrin Beta-1 – Homo Sapiens (Human) | UniProtKB | UniProt Available online: https://www.uniprot.org/uniprotkb/P05556/entry (accessed on 19 September 2024).

71. Harmalkar, A.; Lyskov, S.; Gray, J.J. Reliable Protein–Protein Docking with AlphaFold, Rosetta, and Replica Ex-change. eLife 13, RP94029, doi:10.7554/eLife.94029.

72. Burke, D.F.; Bryant, P.; Barrio-Hernandez, I.; Memon, D.; Pozzati, G.; Shenoy, A.; Zhu, W.; Dunham, A.S.; Albanese, P.; Keller, A.;, et al. Towards a Structurally Resolved Human Protein Interaction Network. Nat. Struct. Mol. Biol. 2023, 30, 216–225, doi:10.1038/s41594-022-00910-8.

73. Chiang, Y.; Hui, W.-H.; Chang, S.-W. Encoding Protein Dynamic Information in Graph Representation for Functional Residue Identification. Cell Rep. Phys. Sci. 2022, 3, 100975, doi:10.1016/j.xcrp.2022.100975.

74. Jiang, Y.; Xu, C.; Zhao, Y.; Ji, Y.; Wang, X.; Liu, Y. LINC00926 Is Involved in Hypoxia-Induced Vascular Endothelial Cell Dysfunction via miR-3194-5p Regulating JAK1/STAT3 Signaling Pathway. Eur. J. Histochem. 2023, 67, doi:10.4081/ejh.2023.3526.

75. He, Q.; He, W.; Dong, H.; Guo, Y.; Yuan, G.; Shi, X.; Wang, D.; Lu, F. Role of Liver Sinusoidal Endothelial Cell in Metabolic Dysfunction-Associated Fatty Liver Disease. Cell Commun. Signal. CCS 2024, 22, 346, doi:10.1186/s12964-024-01720-9.

76. Sgarioto, M.; Vigneron, P.; Patterson, J.; Malherbe, F.; Nagel, M.-D.; Egles, C. Collagen Type I Together with Fibronectin Provide a Better Support for Endothelialization. C. R. Biol. 2012, 335, 520–528, doi:10.1016/j.crvi.2012.07.003.

77. Lindsey, M.L.; Iyer, R.P.; Zamilpa, R.; Yabluchanskiy, A.; DeLeon-Pennell, K.Y.; Hall, M.E.; Kaplan, A.; Zouein, F.A.; Bratton, D.; Flynn, E.R.;, et al. A Novel Collagen Matricryptin Reduces Left Ventricular Dilation Post-Myocardial Infarction by Promoting Scar Formation and Angiogenesis. J. Am. Coll. Cardiol. 2015, 66, 1364–1374, doi:10.1016/j.jacc.2015.07.035.

78. Murdoch, C.; Monk, P.N.; Finn, A. CXC CHEMOKINE RECEPTOR EXPRESSION ON HUMAN ENDOTHELIAL CELLS. Cytokine 1999, 11, 704–712, doi:10.1006/cyto.1998.0465.

79. Emsley, J.; Knight, C.G.; Farndale, R.W.; Barnes, M.J. Structure of the Integrin Α2β1-Binding Collagen Peptide. J. Mol. Biol. 2004, 335, 1019–1028, doi:10.1016/j.jmb.2003.11.030.

80. Emsley, J.; Knight, C.G.; Farndale, R.W.; Barnes, M.J.; Liddington, R.C. Structural Basis of Collagen Recognitionby Integrin Α2β1. Cell 2000, 101, 47–56, doi:10.1016/S0092-8674(00)80622-4.

81. Kamata, T.; Liddington, R.C.; Takada, Y. Interaction between Collagen and the Α2 I-Domain of Integrin Α2β1: CRITICAL ROLE OF CONSERVED RESIDUES IN THE METAL ION-DEPENDENT ADHESION SITE (MIDAS) REGION *. J. Biol. Chem. 1999, 274, 32108–32111, doi:10.1074/jbc.274.45.32108.

82. Smith, C.; Estavillo, D.; Emsley, J.; Bankston, L.A.; Liddington, R.C.; Cruz, M.A. Mapping the Collagen-Binding Site in the I Domain of the Glycoprotein Ia/IIa (Integrin Α2β1) *. J. Biol. Chem. 2000, 275, 4205–4209, doi:10.1074/jbc.275.6.4205.

83. Potel, C.M.; Lemeer, S.; Heck, A.J.R. Phosphopeptide Fragmentation and Site Localization by Mass Spectrometry: An Update. Anal. Chem. 2019, 91, 126–141, doi:10.1021/acs.analchem.8b04746.

84. Martin, D.M.A.; Nett, I.R.E.; Vandermoere, F.; Barber, J.D.; Morrice, N.A.; Ferguson, M.A.J. Prophossi: Automating Expert Validation of Phosphopeptide-Spectrum Matches from Tandem Mass Spectrometry. Bioinforma. Oxf. Engl. 2010, 26, 2153–2159, doi:10.1093/bioinformatics/btq341.

85. Ostrowska-Podhorodecka, Z.; Ding, I.; Lee, W.; Tanic, J.; Abbasi, S.; Arora, P.D.; Liu, R.S.; Patteson, A.E.; Janmey, P.A.; McCulloch, C.A. Vimentin Tunes Cell Migration on Collagen by Controlling Β1 Integrin Activation and Clustering. J. Cell Sci. 2021, 134, jcs254359, doi:10.1242/jcs.254359.

86. Vidyasagar, A.; Wilson, N.A.; Djamali, A. Heat Shock Protein 27 (HSP27): Biomarker of Disease and Therapeutic Target. Fibrogenesis Tissue Repair 2012, 5, 7, doi:10.1186/1755-1536-5-7.

87. Kostenko, S.; Moens, U. Heat Shock Protein 27 Phosphorylation: Kinases, Phosphatases, Functions and Pathology. Cell. Mol. Life Sci. 2009, 66, 3289–3307, doi:10.1007/s00018-009-0086-3.

88. Evans, I.M.; Britton, G.; Zachary, I.C. Vascular Endothelial Growth Factor Induces Heat Shock Protein (HSP) 27 Serine 82 Phosphorylation and Endothelial Tubulogenesis via Protein Kinase D and Independent of P38 Kinase. Cell. Signal. 2008, 20, 1375–1384, doi:10.1016/j.cellsig.2008.03.002.

89. Shah, K.; Shokat, K.M. A Chemical Genetic Screen for Direct V-Src Substrates Reveals Ordered Assembly of a Retrograde Signaling Pathway. Chem. Biol. 2002, 9, 35–47, doi:10.1016/s1074-5521(02)00086-8.

90. Li, Y.; Sun, S.; Zhang, H.; Jing, Y.; Ji, X.; Wan, Q.; Liu, Y. CALU Promotes Lung Adenocarcinoma Progression by Enhancing Cell Proliferation, Migration and Invasion. Respir. Res. 2024, 25, 267, doi:10.1186/s12931-024-02901-3.

91. Chevet, E.; Wong, H.N.; Gerber, D.; Cochet, C.; Fazel, A.; Cameron, P.H.; Gushue, J.N.; Thomas, D.Y.; Bergeron, J.J.M. Phosphorylation by CK2 and MAPK Enhances Calnexin Association with Ribosomes. EMBO J. 1999, 18, 3655–3666, doi:10.1093/emboj/18.13.3655.

92. Augustin-Voss, H.G.; Pauli, B.U. Migrating Endothelial Cells Are Distinctly Hyperglycosylated and Express Specific Migration-Associated Cell Surface Glycoproteins. J. Cell Biol. 1992, 119, 483–491, doi:10.1083/jcb.119.2.483.

93. Worth, D.C.; Daly, C.N.; Geraldo, S.; Oozeer, F.; Gordon-Weeks, P.R. Drebrin Contains a Cryptic F-Actin–Bundling Activity Regulated by Cdk5 Phosphorylation. J. Cell Biol. 2013, 202, 793–806, doi:10.1083/jcb.201303005.

94. Connors, W.L.; Jokinen, J.; White, D.J.; Puranen, J.S.; Kankaanpaöaö, P.; Upla, P.; Tulla, M.; Johnson, M.S.; Heino, J. Two Synergistic Activation Mechanisms of Α2β1 Integrin-Mediated Collagen Binding *. J. Biol. Chem. 2007, 282, 14675–14683, doi:10.1074/jbc.M700759200.

95. Medina-Leyte, D.J.; Domínguez-Pérez, M.; Mercado, I.; Villarreal-Molina, M.T.; Jacobo-Albavera, L. Use of Human Umbilical Vein Endothelial Cells (HUVEC) as a Model to Study Cardiovascular Disease: A Review. Appl. Sci. 2020, 10, 938, doi:10.3390/app10030938.

96. Mischak, H.; Kolch, W.; Aivaliotis, M.; Bouyssié, D.; Court, M.; Dihazi, H.; Dihazi, G.H.; Franke, J.; Garin, J.; de Peredo, A.G.;, et al. Comprehensive Human Urine Standards for Comparability and Standardization in Clinical Proteome Analysis. PROTEOMICS – Clin. Appl. 2010, 4, 464–478, doi:10.1002/prca.200900189.

97. Mokou, M.; Klein, J.; Makridakis, M.; Bitsika, V.; Bascands, J.-L.; Saulnier-Blache, J.S.; Mullen, W.; Sacherer, M.; Zoidakis, J.; Pieske, B.;, et al. Proteomics Based Identification of KDM5 Histone Demethylases Associated with Cardiovascular Disease. EBioMedicine 2019, 41, 91–104, doi:10.1016/j.ebiom.2019.02.040.

98. Megarioti, A.H.; Primo, C.; Kapetanakis, G.C.; Athanasopoulos, A.; Sophianopoulou, V.; André, B.; Gournas, C. The Bul1/2 Alpha-Arrestins Promote Ubiquitylation and Endocytosis of the Can1 Permease upon Cycloheximide-Induced TORC1-Hyperactivation. Int. J. Mol. Sci. 2021, 22, doi:10.3390/ijms221910208.

99. Megarioti, A.H.; Esch, B.M.; Athanasopoulos, A.; Koulouris, D.; Makridakis, M.; Lygirou, V.; Samiotaki, M.; Zoidakis, J.; Sophianopoulou, V.; André, B.;, et al. Ferroptosis-Protective Membrane Domains in Quiescence. Cell Rep. 2023, 42, doi:10.1016/j.celrep.2023.113561.

100. Makridakis, M.; Vlahou, A. GeLC-MS: A Sample Preparation Method for Proteomics Analysis of Minimal Amount of Tissue. Methods Mol. Biol. Clifton NJ 2018, 1788, 165–175, doi:10.1007/7651_2017_76.

101. Claeys, T.; Van Den Bossche, T.; Perez-Riverol, Y.; Gevaert, K.; Vizcaíno, J.A.; Martens, L. lesSDRF Is More: Maximizing the Value of Proteomics Data through Streamlined Metadata Annotation. Nat. Commun. 2023, 14, 6743, doi:10.1038/s41467-023-42543-5.

